# Timed Secreted Proteomes Reveal Regulation of Hepatokines by the Liver Circadian Clock

**DOI:** 10.1101/2025.02.25.640202

**Authors:** Christopher Litwin, Qing Zhang, Ioannis Tsialtas, Zhihong Li, Sophia Hernandez, Steffi Prem, Kristi Dietert, Mallory Keating, Tomoki Sato, Jiyoon Ryu, Lily Q. Dong, Kevin F. Bieniek, Kevin B. Koronowski

## Abstract

Here, we use an ex vivo approach compatible with the circadian timescale to interrogate protein secretion from liver, revealing several findings. Proteomic analyses in male and female mice identify hundreds of proteins that exhibit time-of-day-dependent or clock-dependent secretion involved in extracellular matrix, immune, redox, xenobiotic, and fatty acid functions. Among these, the liver secretes more endostatin, a cleavage product of collagen type XVIII alpha 1 (COL18A1), during the inactive, fasting phase of the diurnal cycle. Temporal regulation of COL18A1/endostatin is dysregulated upon loss of *Bmal1* through combined effects on *Col18a1* transcriptional repression and proteolytic processing. Functional experiments in vivo and in vitro reveal that endostatin suppresses mitochondrial gene expression in white adipose tissue in a time-dependent manner and reduces mitochondrial respiration in adipocytes, while enhancing lipolysis. These results support a mechanism of inter-organ crosstalk whereby hepatically derived, temporally-restricted endostatin tunes adipocytes toward metabolic activities required during the fasting phase.

## INTRODUCTION

Secretion is a primary function of the liver. Grams of albumin are synthesized by the human liver and secreted into the blood each day^1^. The liver supplies the circulation with carrier proteins, apolipoproteins, coagulation factors, complement factors, and hormones^2^, many of which are clinical biomarkers. Proteins secreted by the liver, termed hepatokines, mediate autocrine^3^ and paracrine signaling^4^ within the liver as well as endocrine signaling that underlies inter-organ crosstalk^5^. Hepatokines contribute significantly to the pathophysiology of metabolic disease, and as such, are potential therapeutic targets.

The liver is a temporally dynamic organ, as many of its primary functions exhibit a circadian rhythm. The mouse liver circadian proteome is enriched for well-characterized secreted proteins^6,7^ and a recent study showed that at the transcript and protein level, the clock protein BMAL1 specifically targets and regulates proteins annotated as secreted^8^. In addition, certain hepatokines are regulated by the molecular clock in the liver or mediate circadian functions, including RBP4^9^ and FGF21^10^, two hepatokines implicated in metabolic disease.

Secreted proteins are involved in intercellular and intertissue circadian regulation. An active classical secretory pathway is required for the generation of a robust population-level rhythm among a group of non-neuronal peripheral cells^11^. This cell-cell communication depends on the paracrine signaling action of secreted TGFβ, which aligns the phase of the molecular clock via CRE-mediated immediate early induction of the clock gene *Per2*. Temporal aspects of protein transport from the ER to the plasma membrane are involved in the rhythmic synthesis, assembly, and degradation of collagen, which supports collagen turnover and homeostasis^12^. In addition, the clock repressor Rev-erbα has been identified as a regulator of hepatokines that drive muscle wasting in cancer cachexia^13^. Thus, secretory regulation may be a broad component of circadian control important for homeostasis.

The link between circadian regulation and protein secretion may be especially relevant to endocrine signaling. The liver is quantitatively the largest contributor of protein to the blood and liver-derived proteins are clinical biomarkers. Human studies demonstrate that total protein concentration^14^ and 15-37% of individual proteins exhibit circadian fluctuation in the plasma^15^, with rhythms tied to endogenous circadian control as well as sleeping and eating patterns. Many rhythmic plasma proteins are derived from the liver^16^ and are involved in central systemic processes including hormonal regulation, blood clotting, immune function, molecular transport, energy homeostasis, and xenobiotic metabolism.

Functionally characterizing the full repertoire of liver secreted proteins and their regulatory mechanisms is an active area of study. Hepatic protein secretion comprises the classical secretory pathway^17^, unconventional protein secretion^18^, cargo within small extracellular vesicles^19^, multiple cell types, and specialized anatomical structures^1^. Experimental approaches to study protein secretion include bioinformatic prediction of extracellular localization, proteomic analyses of conditioned media from primary cell cultures^20^ and embedded liver slices^21,22^, and engineered proximity labeling techniques^2,23,24^. Collectively, multiple approaches can facilitate mechanistic insights into hepatokine regulation and function.

Here, we validate an acute, ex vivo assay compatible with the circadian timescale to collect and interrogate proteins secreted from liver. Unbiased mass spectrometry-based proteomic analyses revealed hundreds of time-, clock-, and sex-dependent liver secreted proteins involved in diverse functions. Additional studies showed that secretion of endostatin, a ∼20 kDa cleavage product of the extracellular matrix protein COL18A1, aligned with the inactive, fasting phase and was temporally regulated through combined effects on *Col18a1* transcriptional repression and proteolytic processing. Recombinant endostatin treatment timed to the fasting phase suppressed mitochondrial gene expression in white adipose tissue and subsequent experiments in cultured adipocytes revealed that endostatin lowered mitochondrial respiration and increased lipolysis. Together, these findings indicate a liver-to-adipose signaling axis through which endostatin tunes adipocyte metabolism to promote fasting phase physiology.

## RESULTS

### Protein secretion during acute incubation of liver tissue ex vivo

We reasoned that acute incubation of liver tissue ex vivo would provide an opportunity to capture endogenous circadian regulation from the animal and interrogate protein secretion. We established a workflow compatible with the circadian timescale for acute incubation of ∼100 mg liver tissue, lasting ∼1 h from tissue excision to secreted protein collection (Fig. 1a); ∼100 mg freshly isolated liver was incubated in 5 mL of chemically defined medium continuously bubbled with carbogen for 45 min at 37°C, then the media containing secreted proteins was concentrated ∼20-fold with 3 kDa cutoff centrifugal filters for downstream analyses (see Materials and Methods).

**Fig. 1.**
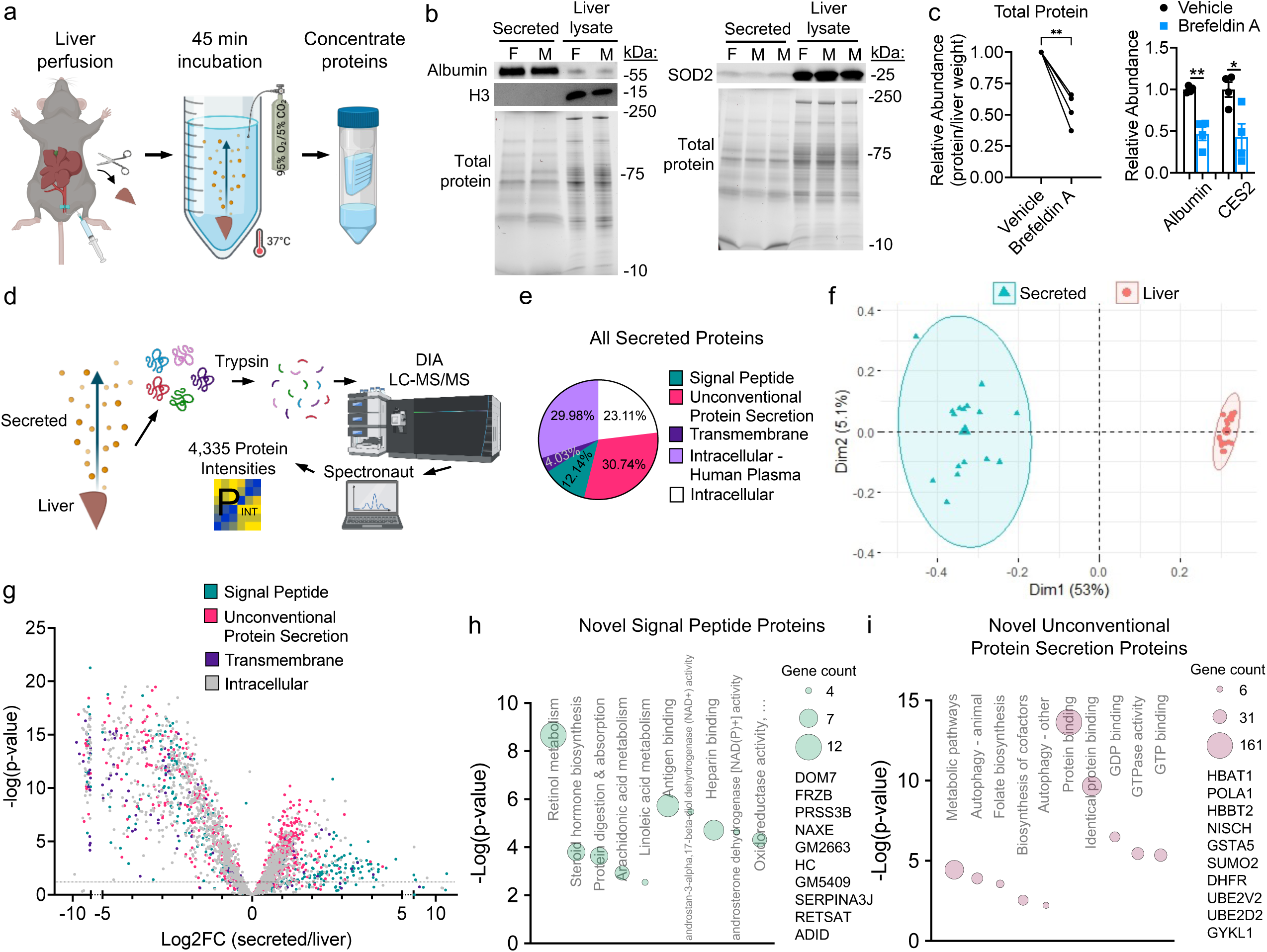
Protein Secretion During Acute Incubation of Liver Tissue Ex Vivo. (a) Simplified scheme of protein secretion from the liver *ex vivo* using C57BL/6J mice at age 12-16 weeks. (b) Western blot from the secreted fraction and liver lysate. F – female; M – male. Total protein from gel. H3 – histone 3; SOD2 – superoxide dismutase 2. (c) Liver was incubated ex vivo with 2 mM Brefeldin A or 2% DMSO vehicle control for 45 min. Left – Two-tailed paired t test of total secreted protein, **p=0.0061, n=4. Right – Western blot quantification of individual secreted proteins, two-tailed paired t test, *p=0.0416, **p=0.0060, n=4. Data are presented as mean +/- SEM. CES2 – carboxylesterase 2. (d) Scheme of proteomic analysis, featuring liver (n=17) and secreted fraction (n=16) from male and female C57BL/6J mice at 12-16 weeks of age. (e) OutCyte classification of all proteins detected in the secreted fraction. Intracellular – Human Plasma = proteins classified as intracellular but identified in the The Human Protein Atlas plasma proteome. (f) Principal component analysis (PCA) of liver and secreted fraction proteomes. (g) Volcano plot showing enrichment of proteins in liver vs secreted fraction. Student’s t-test, two-sided, dashed line p=0.05. Analysis of >5-fold enriched proteins shown in Supplementary Fig. 2g. Proteins are colored by Outcyte classification. (h-i) Gene Ontology enrichment analysis of previously unrecognized secreted proteins. Top 5 pathways (p<0.01, unadjusted) for KEGG and Molecular Function shown. The top 10 most highly abundant proteins are shown to the right. Source data for (b) and (c) are provided in the Source Data file. (a) and (d) Created in BioRender. Koronowski, K. (2026) https://BioRender.com/1e42pue; https://BioRender.com/sf5g35p.

Validation experiments showed that very little IgG was observed in the secreted fraction, indicating negligible blood contamination following perfusion (Supplementary Fig. 1a). Banding patterns of the liver lysate and secreted protein fraction were markedly different, and the secreted fraction was enriched for albumin, negative for the nuclear marker histone 3, and produced a weak signal for the mitochondrial protein SOD2 (Fig. 1b). H&E staining of post-incubation liver sections revealed only ∼2% of cells with fragmented nuclei and ∼5% of cells with cytoplasmic swelling (Supplementary Fig. 1b), indicating minimal cytoplasmic contamination from necrosis or rupture of membranes. Tissue viability was well-maintained, evidenced by no significant drop in liver ATP after 45 min of incubation (Supplementary Fig. 1c) and undetectable levels of the hypoxia-stabilized protein HIF-1α in liver lysates (Supplementary Fig. 1d). Demonstrating dynamic secretion during incubation, the amount of secreted protein increased with incubation time (Supplementary Fig. 1e) and Brefeldin A, an inhibitor of the classical ER-Golgi secretory pathway, reduced protein secretion by ∼50% (Fig. 1c and Supplementary Fig. 1f).

For further validation and identification of liver secreted proteins, we performed DIA LC-MS/MS proteomic analysis on liver and secreted fractions (Fig. 1d). Peptides mapped to 4,197 proteins in liver and 3,135 proteins in the secreted fraction (Supplementary Data 1). Data were consistent among biological replicates, showing Pearson correlation coefficients greater than 0.9 (Supplementary Fig. 2a). Raw intensities spanned 7 orders of magnitude, revealing a large dynamic range of quantification (Supplementary Fig. 2b).

According to OutCyte^25^, nearly half of the secreted proteins contained either a signal peptide (12.14%), a transmembrane domain (4.03%), or fit unconventional protein secretion (30.74%), and the total distribution was similar to published secreted proteomes, which also include a large portion of proteins classified as intracellular^21^ (Fig. 1e, Supplementary Fig. 2c). Notably, greater than 50% of the intracellular-classified proteins are identified in The Human Protein Atlas^26^ plasma proteome, suggesting they may indeed be secreted proteins. Comparing with published liver cell-type-specific proteomes^27–29^, we identified 339 cell-type-specific proteins in the secreted fraction, mapping to hepatocytes (43.95%), Kupffer cells (17.99%), hepatic stellate cells (19.17%), and sinusoidal endothelial cells (18.88%) (Supplementary Fig. 2d).

Principal component analysis (PCA) revealed that liver and secreted fraction samples clustered away from each other along PC1 (53% of variance), indicating substantial differences (Fig. 1f). As expected, the secreted fraction contained well characterized secreted proteins and was enriched for extracellular compartments and systemic functions including immunoglobulin production, complement activation, and innate immune response, whereas the liver featured highly abundant metabolic enzymes and was enriched for intracellular compartments and mitochondrial functions (Supplementary Fig. 2e, Supplementary Data 2).

We analyzed relative abundance between the liver and secreted fraction, reasoning that bona fide secreted proteins will be more abundant in the secreted fraction. Indeed, individual plasma proteins (albumin, >27-fold; hemopexin, >22-fold; transferrin, >9-fold) and secreted protein families (serpins, immunoglobulins, apolipoproteins, and complement factors, ∼5-fold) were more abundant in the secreted fraction (Supplementary Fig. 2f). Considering proteins more than 5-fold more abundant, 67.68% contained N-terminal signal peptides (Fig. 1g and Supplementary Fig. 2f). Conversely, proteins that were >5-fold more abundant in the liver tended to be intracellular-designated proteins (50.3%) and were associated with intracellular functions, especially mitochondrial processes (Supplementary Fig. 2g, Supplementary Data 2).

Notably, the secreted fraction contained more than 500 signal peptide-containing or unconventionally secreted proteins not identified in 7 previously published secreted proteomes^2,20,22–24,30,31^, which were involved in known as well as previously unappreciated secretory functions of the liver (Fig. 1h-i, Supplementary Fig. 2h). These data show that an acute ex vivo approach is suitable to interrogate hepatic protein secretion.

### Identification of time-of-day-dependent liver secreted proteins

Next, we used the acute ex vivo approach to identify time-of-day-dependent secreted proteins. Liver tissue from male and female mice was excised during the inactive, fasting phase at zeitgeber time (ZT) 8 (8 h after lights on) or active, feeding phase at ZT20 (8 h after lights off) (n=4 per sex, per time point). These timepoints reflect opposite phases of the molecular clock machinery and align with previous reports on the temporal accumulation of secreted proteins within the liver^6,7^. Analyses with and without sex as a variable revealed 60 time-dependent secreted proteins (Fig. 2a-b), several of which had known ties to circadian rhythm (FGB and FBLN1^32^, ACOT1, 3, and 4^33^, and LGALS1^34^). Several hits were further validated as secreted proteins in conditioned media from primary hepatocyte cultures (Supplementary Fig. 3a).

**Fig. 2.**
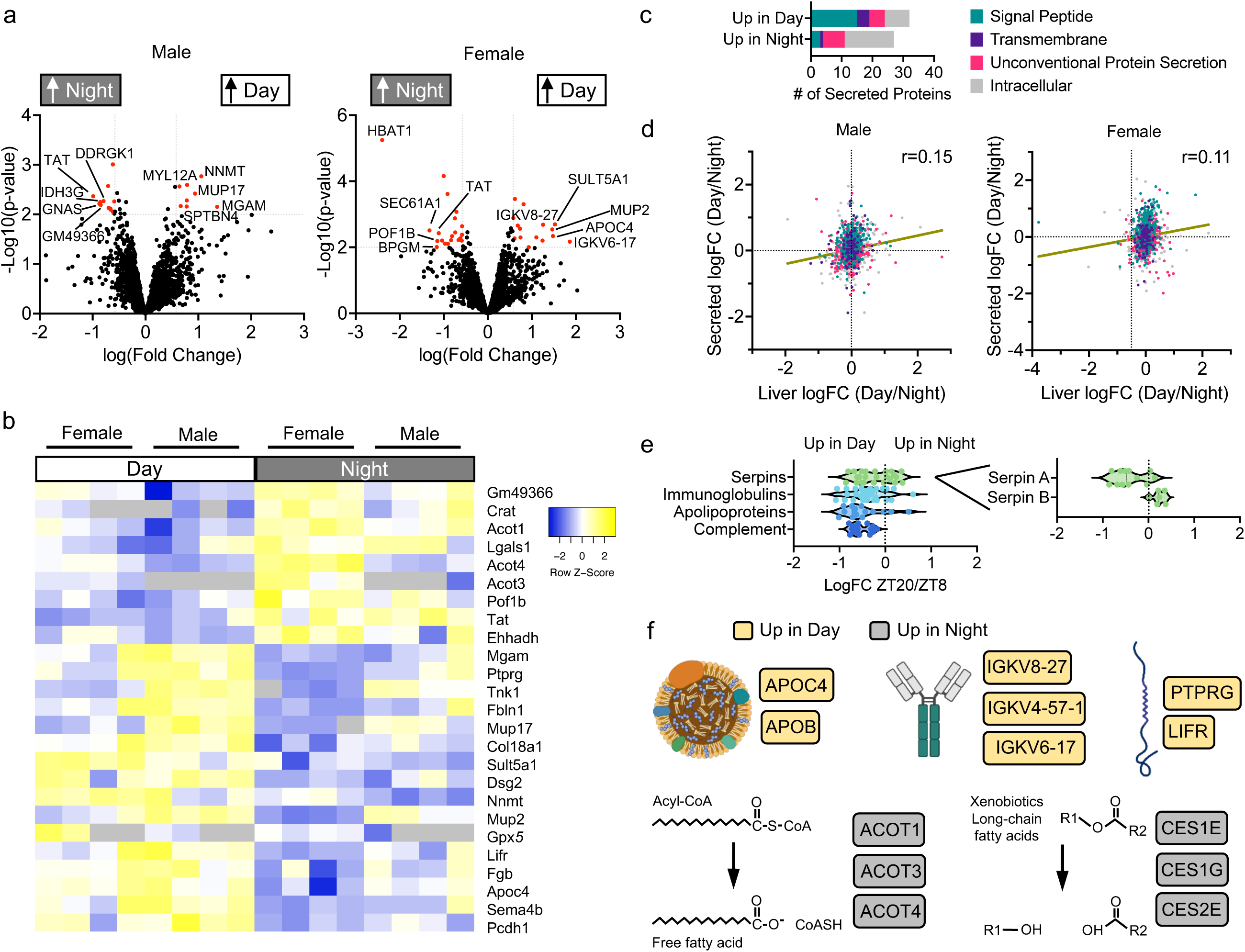
Time-of-Day Defines a Subset of Liver Secreted Proteins. Analysis of secreted fraction proteomes from two diurnal time points, ZT8 (inactive, fasting phase) and ZT20 (active, feeding phase), for male and female C57BL/6J mice at 12-16 weeks of age (n=4 per sex, per group). (a) Volcano plots showing time-dependent secreted proteins. Red data points = Student’s t-test, two-sided, p<0.01 (unadjusted), > ± 50% change. The top 5 proteins by logFC are labeled. (b) Heatmap of secreted proteins exhibiting time dependence as defined in (a) from combined sexes analysis. (c) OutCyte classifications for all time-dependent secreted proteins. (d) Plot of logFC (ZT8/ZT20) in the liver and secreted fraction for all proteins quantified in both compartments. Pearson correlation (r), simple linear regression line shown. (e) Fold-Change analysis of classical plasma protein families. Clade A and B serpins, which exhibit differential time-dependent secretion, are shown separately to the right. (f) Example proteins with time-dependent secretion. APOC4 – apolipoprotein C-IV; APOB – apolipoprotein B; IGKV8-27 – immunoglobulin kappa chain variable 8-27; IGKV4-57-1 – immunoglobulin kappa variable 4-57-1; IGKV6- 17 – immunoglobulin kappa variable 6-17; PTPRG – protein tyrosine phosphatase receptor type G; LIFR – leukemia inhibitory factor receptor; ACOT – acyl-CoA thioesterase; CES – carboxylesterase. Created in BioRender. Koronowski, K. (2026) https://BioRender.com/mqqot8y.

While the number of proteins secreted more in day or night was similar, proteins secreted more in day had more well-defined secretory regulation, with 46.88% containing a signal peptide (Fig. 2c). Proteins secreted more in night favored unconventional protein secretion or were labeled as intracellular. There was a very weak positive correlation between log fold-changes (ZT8/ZT20) in the liver and secreted fraction (male, r=0.15; female, r=0.11), indicating that, in general, liver protein abundance does not primarily dictate time-dependent secretion (Fig. 2d).

Many proteins secreted more in day belonged to well characterized plasma protein families. We plotted log fold-changes for all quantified immunoglobulins, apolipoproteins, serpins, and complement factors and found that their secretion was increased at ZT8 by ∼35% on average (Fig. 2e). Most apolipoproteins tended to be secreted more at ZT8 (exceptions: APOA5, APOL7C, and APOL9B), and APOB and APOC4 were highly significant (Fig. 2f). An abnormally high blood APOB value is a clinical indicator of cardiovascular disease^35^, and by comparison, little is known about the function of APOC4. In addition, two receptors – leukemia inhibitory factor receptor (LIFR) and protein tyrosine phosphatase receptor type G (PTPRG) – were increased in the secreted fraction at ZT8 (Fig. 2f). Transmembrane receptors are known to undergo proteolytic cleavage, also referred to as ectodomain shedding, which releases the extracellular fragment of the protein to circulate as a soluble signaling peptide. COL18A1/endostatin was a notable protein secreted more in day.

Among the proteins displaying increased secretion at ZT20 were several enzymes related to lipid metabolism including CES1, CES2, ACOTs, ACOX1, EHHADH, ACP6, FDX1, and CRAT (Fig. 2f). Carboxylesterases (CESs) participate in lipid metabolism by hydrolyzing esters and thioesters intracellularly and in the circulation. CES2 was recently shown to be secreted by the liver in response to increasing lactate concentrations during exercise and, in its secreted form, protect against high fat diet induced obesity^24^. The other enzymes identified here may participate in circadian regulation of metabolism as secreted proteins, yet future studies are needed to further validate secretion and secretory functions.

An analysis of sex-dependent proteins revealed 176 male-only and 147 female-only secreted proteins (Supplementary Fig. 3b, Supplementary Data 1), 74 secreted proteins showing a sex difference at both ZT8 and ZT20, as well as 167 ZT8 sex-dependent proteins and 180 ZT20 sex-dependent proteins (Supplementary Fig. 3c-d). There was a moderate positive correlation between log fold-changes (male/female) in the liver and secreted fraction (ZT8, r=0.46; ZT20, r=0.50), indicating that liver protein abundance may dictate sex-dependent secretion to a certain degree (Supplementary Fig. 3e). Lipid metabolic process and sulfation were the top enrichments of proteins secreted more in females and included several well-characterized intracellular enzymes of lipid metabolism with unknown but potentially important secreted functions given that they circulate in human plasma (MVD, AKR1D1, SCLY, HADH), as well as four sulfotransferases^36^ (SULT1A1, SULT1C2, SULT1D1, SULT3A1) (Supplementary Fig. 3f). Proteins secreted less in females included other xenobiotic enzymes, complement activation factors C8A and C8B, and 5 serine protease inhibitors (Serpins; SERPINA1A, SERPINA1E, SERPINA3J, SERPINA3K, SERPINB1A) (Supplementary Fig. 3f). Other notable sex-dependent secreted proteins were selenium-binding protein 2 (SELENBP2), apolipoprotein E (APOE), haptoglobin (HP), gamma-glutamyltransferase 1 (GGT1) and CES3.

### The hepatocyte clock regulates hepatic protein secretion

Time-dependent secretion may involve temporal regulation stemming from the molecular clock. We measured hepatic mRNAs for a set of ten time-dependent secreted proteins and found extensive differential expression in hepatocyte-specific *Bmal1* knockout mice (*Bmal1*^hep-/-^) (Supplementary Fig. 4a). Altered expression was appreciable at the protein level; for example, FGB was upregulated at several diurnal timepoints, especially during the inactive, fasting phase (Supplementary Fig. 4b,c). Luciferase reporter assays in AML12 cells revealed that BMAL1 and CLOCK transactivate the *Fgb* promoter, linking clock-based transcriptional regulation to secreted protein expression (Supplementary Fig. 5a). In addition, we measured responses to circadian misalignment and found that chronic jet lag shifted or deregulated the expression of secreted proteins (Supplementary Fig. 4d-h), including 6 secreted proteins identified herein and 8 robustly circadian genes encoding signal peptide-containing proteins identified previously^37^. These data provided further impetus to determine the contribution of the molecular clock to hepatic protein secretion.

We performed ZT8 vs ZT20 ex vivo secretion assays using liver from wild-type (WT) and *Bmal1*^hep-/-^ mice (n=4 per sex, per time point). Proteomic analyses revealed that most of the WT time-dependent secreted proteins (82 proteins, including 15 male specific and 38 female specific) lost time-dependence in *Bmal1*^hep-/-^ mice (83.7%, 74.4%, and 90.9% from combined, male, and female analyses, respectively) (Fig. 3a, Supplementary Fig. 6a, Supplementary Data 3), indicating that the hepatocyte clock is involved in temporal secretion. Among these proteins were the extracellular matrix components COL18A1/endostatin, LUM, and EFEMP1, extracellular superoxide dismutase (SOD3), and IGF binding protein 2 (IGFBP2), which exhibited increased secretion at ZT8 (Fig. 3b). Conversely, many proteins involved in the immune response were secreted more at ZT20, including IGKV family immunoglobulins, IGHG2C, and IGH-3. Several proteins involved in carbohydrate metabolism lost temporal secretion in *Bmal1*^hep-/-^ (i.e., MGAM, LANCL2, AMY1, GUSB). Separate male and female analyses revealed significant enrichments (Supplementary Fig. 6b, Supplementary Data 2). Loss of hepatocyte *Bmal1* also resulted in *de novo* time-dependent secretion of proteins (Fig 3a, Supplementary Fig. 6a,c, Supplementary Data 2), an effect that was more pronounced in males.

**Fig. 3.**
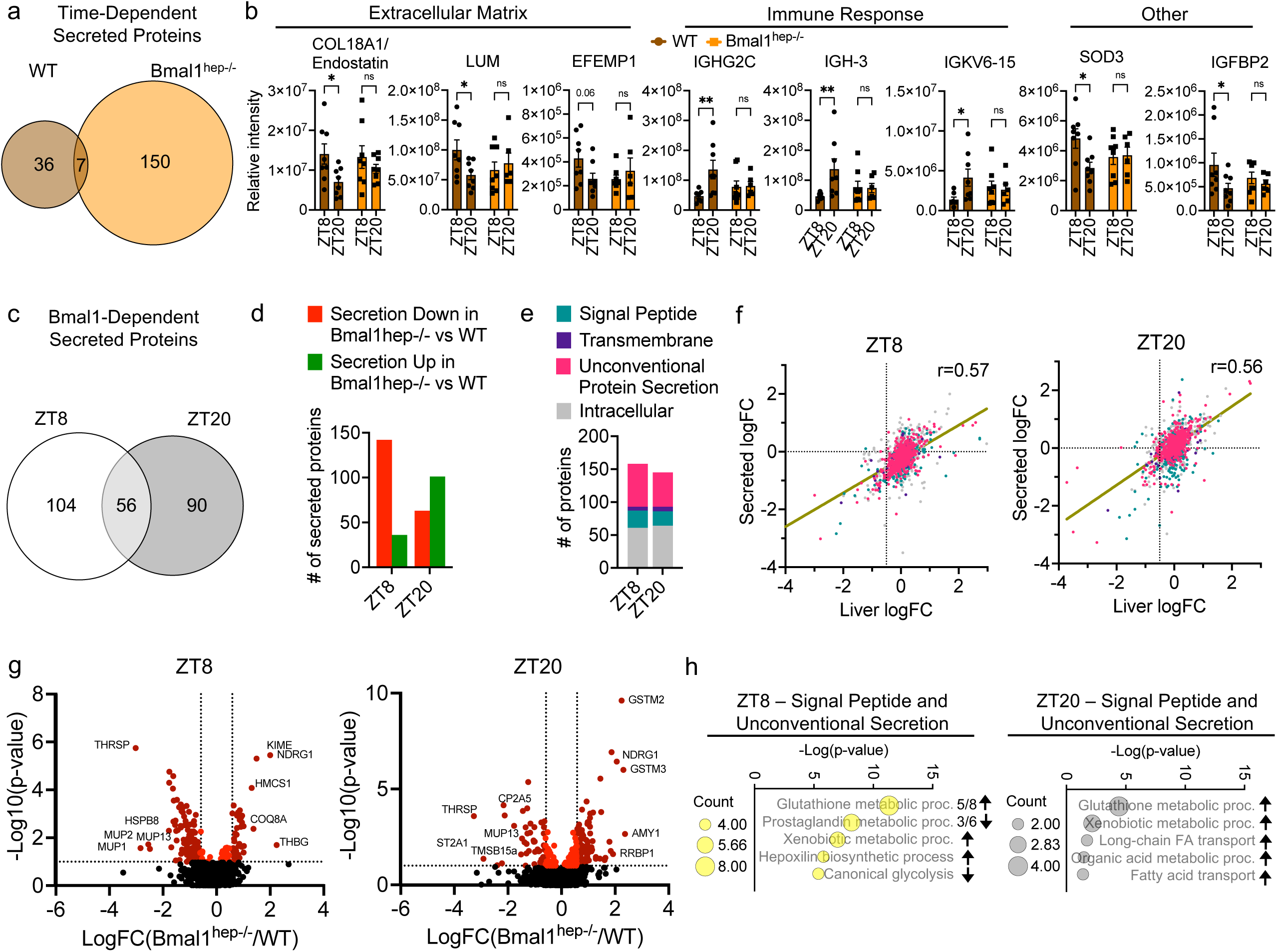
Hepatocyte *Bmal1* Regulates Hepatic Protein Secretion. Proteomic analyses of liver ex vivo secreted fractions of WT and hepatocyte-specific *Bmal1* knockout (*Bmal1*^hep-/-^) male and female mice aged 12-16 weeks at two diurnal time points, ZT8 (inactive, fasting phase) and ZT20 (active, feeding phase), n=4 per sex, per group. (a) Comparison of time-dependent secreted proteins in each genotype, Student’s t-test, two-sided, FDR <0.1, > ± 50% change. (b) Example WT time-dependent secreted proteins that lost time-dependence in *Bmal1*^hep-/-^, Two-way ANOVA, Fisher’s LSD, COL18A1 *p=0.0313, LUM *p=0.0369, IGHG2C **p=0.0069, IGH-3 **p=0.0089, IGKV6-15 *p=0.0236, SOD3 *p=0.0119, IGFBP2 *p=0.0342. Data are presented as mean +/- SEM. Outliers removed by Grubb’s test (p<0.05). COL18A1 – collagen type XVIII alpha 1 chain; LUM – lumican; EFEMP1 – EGF containing fibulin extracellular matrix protein 1; immunoglobulin heavy constant gamma 2C; immunoglobulin heavy constant gamma 2B/heavy chain 3; immunoglobulin kappa variable 6-15; SOD3 – superoxide dismutase 3; IGFBP2 – insulin-like growth factor binding protein 2. (c-e) Comparison of *Bmal1*-dependent secreted proteins at each time point, FDR <0.1, > ± 50% change. Proteins without identifiers removed from analysis. (d) Directionality of *Bmal1*-driven secretory effects. (e) Outcyte classification of secretory mechanism. (f) Same color coding as in (e). Plot of logFC (*Bmal1*^hep-/-^/WT) in the liver and secreted fraction for all proteins quantified in both compartments. Pearson correlation (r), simple linear regression line shown. (g) Volcano plots showing individual *Bmal1*-dependent secreted proteins. Dark red data points = Student’s t-test, two-sided, FDR <0.1, > ± 50% change. The top 5 proteins by logFC are labeled. (h) Gene Ontology enrichment analysis (Biological Process) of *Bmal1*-dependent secreted proteins that contain signal peptides or are unconventionally secreted. The arrows indicate the direction of the change. ZT – zeitgeber time.

Comparing genotypes at ZT8 and ZT20 revealed 250 *Bmal1*-dependent secreted proteins, with 104 unique to ZT8 and 90 unique to ZT20 (Fig. 3c). An additional 33 male specific and 43 female specific *Bmal1*-dependent secreted proteins were identified (Supplementary Fig. 6d-e). Secretion of *Bmal1*-dependent proteins tended to be increased at ZT8 (Fig. 3d), and total serum protein was typically steady across the diurnal cycle yet dipped at ZT8 in *Bmal1*^hep-/-^ mice or under chronic jet lag (Supplementary Fig. 5b)., indicating that the clock promotes secretion during the inactive, fasting phase. *Bmal1*-dependent secreted proteins varied in terms of secretory mechanism (Fig. 3e), and there was a moderate positive correlation between log fold-changes (*Bmal1*^hep-/-^/WT) in the liver and secreted fraction (ZT8, r=0.57; ZT20, r=0.56), showing that the effect of *Bmal1* on the liver proteome influences the secretome (Fig. 3f).

Volcano plots show the significant *Bmal1*-dependent secreted proteins at ZT8 and ZT20 (Fig. 3g). Gene ontology enrichment analysis of signal peptide containing and unconventionally secreted proteins revealed glutathione metabolism as a top *Bmal1*-dependent secretory function (Fig. 3h, Supplementary Data 2). Seven glutathione S-transferases (GSTs) of the mu class were secreted more in *Bmal1*^hep-/-^, whereas alpha GSTs and theta GSTs were secreted less. Enzyme families involved in detoxification of xenobiotics were also identified, with 3 sulfotransferases (SULT1A1, SULT2A5, SULT2A1) exhibiting increased secretion, and 3 cytochrome P450 proteins (CYP2C68, CYP2C29, CYP2C39) exhibiting decreased secretion, in *Bmal1*^hep-/-^. Notably, sulfation was a top sex-dependent secretory pathway (xenobiotic enrichment terms). Hepatic lipase (LIPC) and fatty acid binding proteins 1 and 4 (FABP1, FABP4) were secreted more in *Bmal1*^hep-/-^, especially during the active/feeding phase at ZT20, and apolipoprotein a 4 (APOA4) and several carboxylesterases (CES1C, CES1D, CES1F) were secreted less in *Bmal1*^hep-/-^. These results show that the hepatocyte clock regulates protein secretion tied to hepatic and systemic functions.

### The hepatocyte clock regulates *Col18a1* and endostatin

COL18A1 was a notable time- and clock-dependent secreted protein. COL18A1 is an extracellular matrix component of collagen that is proteolytically cleaved at its C-terminus to produce endostatin, a soluble ∼20 kDa peptide. Recent rodent studies report beneficial metabolic effects of chronic recombinant endostatin treatment^38,39^, yet the physiological sources, secretion mechanisms, diurnal activities of endostatin remain poorly defined.

Across two independent datasets, livers from WT mice secreted more COL18A1/endostatin at ZT8 during the inactive, fasting phase as compared to ZT20 during the active, feeding phase, whereas livers from *Bmal1*^hep-/-^ mice showed no time-of-day difference in secretion, which was similarly elevated at ZT8 and ZT20 (Fig. 3). To determine the involvement of rhythmic transcription, we probed hepatic mRNA and found that *Col18a1* displayed a small amplitude rhythm in WT liver, peaking near ZT4 (Fig. 4a). This rhythm was lost in *Bmal1*^hep-/-^ liver, wherein *Col18a1* was consistently upregulated across many diurnal time points. Specific transcripts resulting from alternative polyadenylation may also contribute to rhythmic accumulation of *Col18a1* mRNA (Greenwell et al., BioRxiv, 2020).

**Fig. 4.**
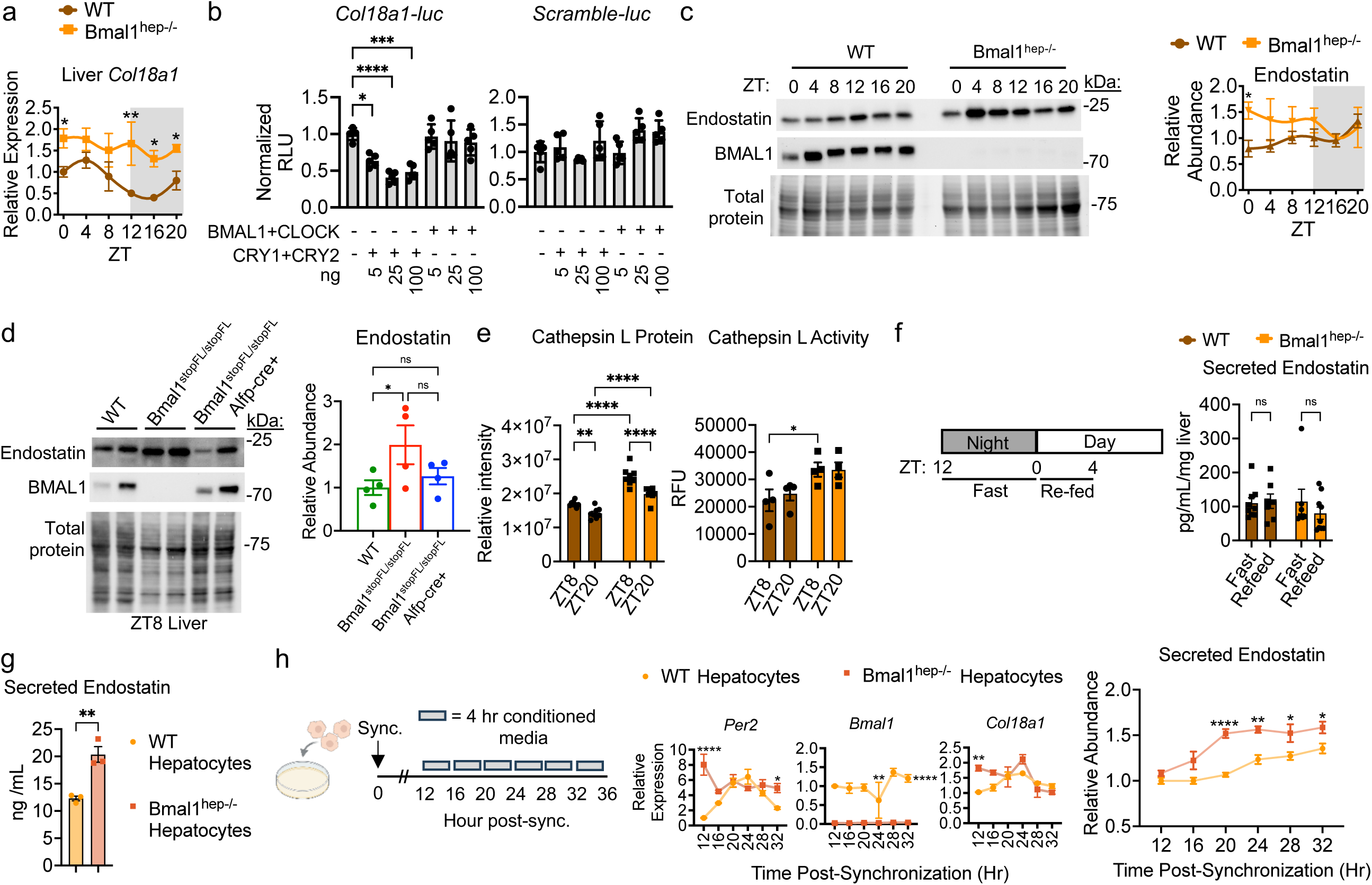
Hepatocyte Clock Proteins Regulate *Col18a1* Transcription and Endostatin Secretion. (a-h) Data are presented as mean +/- SEM. (a) Expression of *Col18a1* by qPCR in the liver at 6 diurnal timepoints. Two-way ANOVA, Fisher’s LSD post hoc test, ZT0 *p=0.0399, ZT12 **p<0.0033, ZT16 *p=0.0112, ZT20 *p=0.0335. WT ZT0 and *Bmal1*^hep-/-^ ZT12 n=3, all other groups n=4 mouse livers. Gray shading indicates lights off. Rhythmicity analyses – Cosinor: WT p=0.00993, *Bmal1*^hep-/-^ p=0.8212; RAIN: WT p=0.083, *Bmal1*^hep-/-^ p=0.985. (b) Transient transfection assay in AML12 hepatocytes. Luciferase driven by *Col18a1* promoter region (-2 kb to + 1kb relative to TSS). Scramble-luc = randomized DNA sequence control. One-way ANOVA, Tukey’s post hoc test, *p=0.0132, ***p=0.0002, ****p=<0.0001, n=5. One representative experiment shown. See also Supplementary Fig. 7a. (c-d) Western blot of whole liver lysates. (c) Two-way ANOVA, Fisher’s LSD post hoc test, *p<0.05. WT ZT4, 8, 16 and *Bmal1*^hep-/-^ ZT4, 12, 20 n=3, all other groups n=4 mouse livers. Rhythmicity analyses – Cosinor: WT p=0.6118, *Bmal1*^hep-/-^ p=0.2977; RAIN: WT p=0.99, *Bmal1*^hep-/-^ p=0.38. (d) One-way ANOVA, Fisher’s LSD post hoc test, *p=0.0438, n=4 mouse livers. (e) Left – Cathepsin L protein abundance in liver from proteomics dataset, n=8. Right – Cathepsin L enzymatic activity from liver, n=4. Left and Right – Two-way ANOVA, Fisher’s LSD post hoc test, *p<0.05, **p<0.01, ***p<0.001, ****p<0.0001. (f) Acute ex vivo secretion assay on livers of fasted or fasted-refed mice. Secreted endostatin measured by ELISA. One-way ANOVA, Fisher’s LSD post hoc test, WT refeed male and *Bmal1*^hep-/-^ fast male n=3, WT refeed female, WT fast female, *Bmal1*^hep-/-^ refeed female, and *Bmal1*^hep-/-^ fast female n=4, WT fast male and *Bmal1*^hep-/-^ refeed male n=5. (g) 4-h conditioned media generated from unsynchronized primary hepatocytes. Secreted endostatin measured by ELISA. Student’s t-test, two-sided, **p=0.0056, n=3. One representative experiment shown. (h) Left – schematic of primary hepatocyte synchronization with dexamethasone (1 h, 100 nM) and subsequent collection of conditioned media from separate wells at 4 h intervals. Created in BioRender. Koronowski, K. (2026) https://BioRender.com/zg2obsf. Middle – qPCR confirming synchronization. Two-way ANOVA, Sidak’s post hoc test, *Bmal1* **p=0.0096, ****p=<0.0001, *Per2* *p=0.0353, ****p=<0.0001, *Col18a1* **p=0.0055. All groups for *Per2* and *Col18a1* n=3 except for WT Hr 24 n=2. All groups for *Bmal1* n=3 except for WT Hr 24 and *Bmal1*^hep-/-^ Hr 20 n=2. Right – Quantification of secreted endostatin by ELISA. Two-way ANOVA, Sidak’s post hoc test, *p=0.0251 (Hr 28), *p=0.0459 (Hr 32), **p=0.022, ****p=<0.0001, n=3. One representative experiment shown. Source data are provided in the Source Data file. ZT – zeitgeber time.

We analyzed the *Col18a1* promoter and identified several putative E-box motifs. In addition, published cistromes indicate that the clock repressor proteins CRY1 and CRY2 bind near the *Col18a1* promoter at sites that do not overlap with putative E-boxes, and *Col18a1* is upregulated in the liver of *Cry1*^-/-^;*Cry2*^-/-^ mice^40^ (Supplementary Fig. 7a,b). To directly test transcriptional regulation by clock proteins, we generated a luciferase reporter construct with a 3 kb sequence of the *Col18a1* promoter (-2 kb to +1kb relative to TSS; *Col18a1*-luc). In transient transfection assays using AML12 cells, CLOCK and BMAL1 failed to activate *Col18a1*-luc, yet CRY repressed *Col18a1*-luc by ∼50% (Fig. 4b). CRY did not repress a scrambled DNA control luciferase reporter. These findings suggest that the repressive arm of the clock, or CRY proteins in concert with other transcription factors, temporally regulate *Col18a1* transcription.

At the protein level, endostatin was upregulated during the early inactive, fasting phase in the liver of *Bmal1*^hep-/-^ mice (Fig. 4c) and whole-body *Bmal1* knockout mice (*Bmal1*^stopFL/stopFL^) (Fig. 4d). In mice expressing *Bmal1* only in hepatocytes and in no other cell types (*Bmal1*^stopFL/stopFL^;Alfp-Cre^tg/0^)^41^, liver endostatin was restored, at least in part, to WT abundance (Fig. 4d). An increase in endostatin could also stem from altered activities of enzymes that cleave COL18A1, such as cathepsin L, a cysteine proteinase involved in extracellular matrix remodeling^42^. Cathepsin L protein abundance and enzymatic activity were upregulated in *Bmal1*^hep-/-^ liver, especially at ZT8 (Fig. 4e), which coincides with the observed increases in endostatin abundance and secretion.

Nutrient availability is known to regulate hepatic rhythmicity and protein secretion. To determine if feeding or fasting regulates endostatin secretion, we performed the ex vivo secretion assay on livers from mice fasted for 12 h from ZT12 to ZT24/0 or fasted and refed for 4 h from ZT24/0 to ZT4. Fasting increased liver *Col18a1* expression by ∼30% in WT and *Bmal1*^hep-/-^ mice (Supplementary Fig. 7c), yet fasted liver and refed liver secreted a similar amount of endostatin, irrespective of genotype (Fig. 4f). These data indicate that the hepatocyte clock drives temporal regulation of endostatin secretion. To test this, we first confirmed that endostatin is secreted from primary hepatocytes, and that hepatocytes obtained from *Bmal1*^hep-/-^ mice secrete more endostatin, consistent with ex vivo data (Fig. 4g). Then, we synchronized hepatocytes with dexamethasone and incubated them with serum-free media at 4 h intervals starting 12 h post-synchronization until 36 h. Clock gene expression profiles confirmed efficient synchronization, and we found that endostatin secretion peaked earlier in *Bmal1*^hep-/-^ hepatocytes and was increased at several timepoints compared to WT hepatocytes (Fig. 4h). Altogether, these results indicate that temporal secretion of endostatin stems from a hepatocyte-clock-autonomous mechanism involving transcriptional repression of *Col18a1* and cleavage of COL18A1.

### Endostatin tunes adipocyte metabolism to the inactive, fasting phase

Next, we sought to determine if hepatocyte *Bmal1* influences circulating abundance of endostatin which could, in turn, affect distal tissues via endocrine signaling. In WT mice, serum endostatin was time-of-day-dependent, with a small peak at ZT8 (Fig. 5a). This peak was not observed in *Bmal1*^hep-/-^ mice, which exhibited elevated serum endostatin at the dark to light transition (ZT0).

**Fig. 5.**
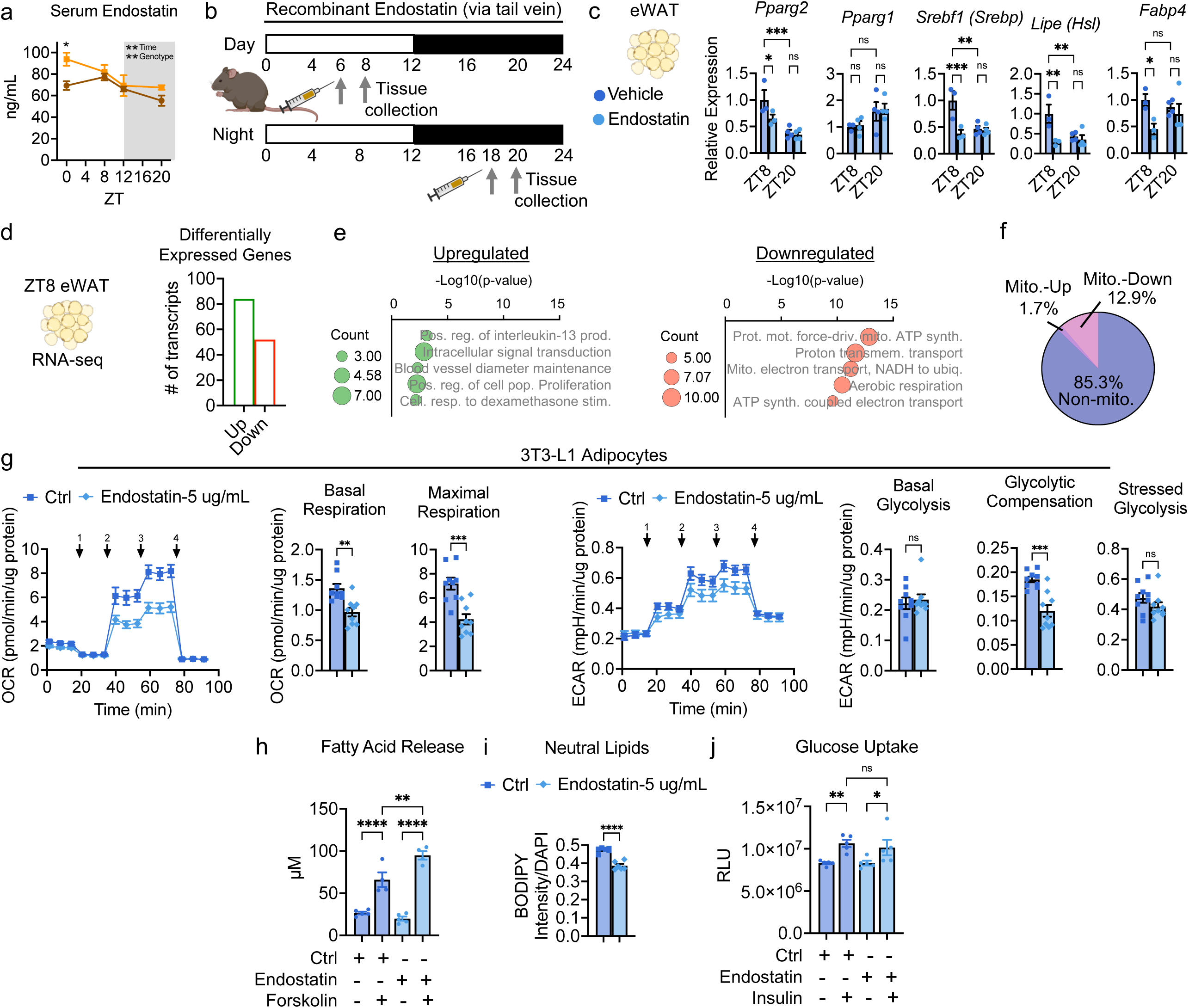
Endostatin Tunes Adipocyte Metabolism During the Fasting Phase of the Diurnal Cycle. (a) Serum endostatin measured by ELISA from adult male mice at the indicated timepoints, Two-way ANOVA, Sidak’s post hoc test, time **p=0.0025, genotype **p=0.0076, ZT0 *p=0.0127. All groups n=5 except for *Bmal1*^hep-/-^ ZT20 n=4. Gray shading indicates lights off. Data are presented as mean +/- SEM. (b) Scheme of experimental design for timed injection of 100 ug recombinant human endostatin into 15-week-old C57BL/6J male and female mice. (c) qPCR in epididymal white adipose tissue (eWAT) from female mice. Two-way ANOVA, Fisher’s LSD post hoc test, *Pparg2* *p=0.0320, ***p=0.0009, *Srebf1* ***p=0.0010, **p=0.0017, *Lipe* ZT8 **p=0.0028, ZT8 vs ZT20 **p=0.0076, *Fabp4* *p=0.0252, n=3-4 (n-value for each group shown within graph). Data are presented as mean +/- SEM. See also Supplementary Fig. 8. (d-f) Total RNA-sequencing of eWAT from females at ZT8, n=4. (d) Directionality of differentially expressed genes (DEGs), FDR <0.05, > ± 50% change. (e) Gene Ontology enrichment analysis (Biological Process) of up or down regulated genes, p<0.01 unadjusted. (f) Specific analysis of mitochondrial DEGs from MitoCarta 3.0. (g-j) Experiments in differentiated 3T3-L1 adipocytes. One representative experiment shown. Data are presented as mean +/- SEM. (g) Seahorse assay. Treatment with recombinant human endostatin for 24 hr prior. Arrows indicate application of (1) 1.0 uM oligomycin, (2) 1.0 uM FCCP, (3) 1.0 uM FCCP, (4) 0.5 uM rotenone/antimycin A. OCR = oxygen consumption rate; ECAR = extracellular acidification rate. Student’s t-test, two-sided, OCR basal **p=0.0014, OCR maximal ***p=0.0004, ECAR compensation ***p=0.0002, n=9. (h) Measurement of non-esterified fatty acids in the media following 2-h forskolin (5 uM) and/or 5 ug/mL endostatin treatment. Two-way ANOVA, Fisher’s LSD post hoc test, **p=0.0010, Ctrl ****p=<0.0001, Endostatin ****p=<0.0001. Without forskolin groups n=5, with n=4. (i) BODIPY 493/503 intensity normalized to cell number. Endostatin treatment for 24 h prior, Student’s t-test, two-sided, ****p=<0.0001, n=6. (j) 2-deoxyglucose uptake assay. Insulin (100 nM) treatment for 1 h. 5 ug/mL endostatin treatment for 2 h prior to insulin incubation and during incubation. Two-way ANOVA, Fisher’s LSD post hoc test, **p=0.0061, *p=0.0254, n=5. Source data for (a), (c), and (g)-(j) are provided in the Source Data file. ZT – zeitgeber time. (b-d) Created in BioRender. Koronowski, K. (2026) https://BioRender.com/byyuqa5.

To interrogate time-dependent endocrine effects of endostatin, adult 15-week-old male and female mice were administered tail vein injections of 100 ug recombinant human endostatin or vehicle control at ZT6 or ZT18, near the natural peak and trough of serum endostatin, respectively (Fig. 5b). Based on the ∼3 h half-life of recombinant endostatin^43^, tissue was collected 2 h post-injection. Food was removed during this 2 hour period to control for acute food intake. Blood glucose was not significantly altered by vehicle or endostatin compared to pre-injection values (Supplementary Fig. 8a,b). Analysis of epididymal white adipose tissue (eWAT), a previously identified target of endostatin, revealed that endostatin lowered the expression of several genes that are critical to the metabolic activities of adipocytes, including the *Pparg2* isoform that regulates adipogenesis^44^ (*Pparg2*, *Srebf1*, *Lipe*, *Mgll*, *Fabp4*). These effects were exclusive to ZT6 injection and female mice (Fig. 5c, Supplementary Fig. 8c).

To determine transcriptome-wide effects of endostatin, we performed bulk RNA sequencing (RNA-seq) on eWAT collected at ZT8 from female mice (n=4). We identified 136 differentially expressed genes, including genes identified by qPCR (Fig. 5d). Gene Ontology enrichment analysis revealed that all of the top 5 pathways downregulated by endostatin were involved in mitochondrial metabolism (Fig. 5e, Supplementary Data 2). According to MitoCarta3.0 (ref), a large portion of endostatin-modulated genes (∼15%) were mitochondrial genes, and nearly all were downregulated (Fig. 5f).

To evaluate the functional implications of the RNA-seq results, we differentiated 3T3-L1 adipocytes in vitro and probed their metabolism. Consistent with downregulation of mitochondrial genes, 5 ug/mL endostatin treatment lowered the oxygen consumption rate under basal conditions and especially when adipocytes were metabolically stressed (Fig. 5g). Endostatin also lowered the extracellular acidification rate, indicative of a change in glycolytic rate. Impaired adipocyte differentiation is shown to coincide with reduced mitochondrial respiration^45^. The observation that endostatin lowers mitochondrial respiration is thus consistent with previous reports suggesting that endostatin inhibits adipogenesis^39^. Suppression of adipogenesis occurs alongside increased lipolysis during the fasting phase of the diurnal cycle. Given that the observed effects of endostatin were exclusive to the fasting phase (ZT8), we tested whether endostatin regulates lipolysis. An acute 2-hr exposure to endostatin enhanced forskolin-induced fatty acid release from adipocytes (Fig. 5h) and, evidenced by BOIDPY staining, endostatin treatment reduced adipocyte neutral lipid stores (Fig. 5i). Conversely, endostatin had no effect on insulin-stimulated glucose uptake (Fig. 5j), which fuels *de novo* lipogenesis and increases lipid storage in adipocytes during feeding. Together, these findings support an endostatin-mediated liver-to-WAT signaling axis that enhances lipolysis during the inactive, fasting phase of the diurnal cycle.

## DISCUSSION

In this study, we identified hepatokines regulated by time-of-day, the hepatocyte clock, and sex, involved in various functions. We further characterized the temporal secretion mechanism and temporal functional effect of one prominent hepatokine, COL18A1/endostatin, finding that heightened fasting phase secretion of endostatin acts on adipocytes to lower mitochondrial respiration and promote lipolysis. Altogether, our results implicate the temporal regulation of hepatokines in systemic homeostasis.

Previous studies show that secreted proteins are enriched in the liver circadian proteome at specific times of day^6,7^. Proteins comprising the secretory machinery in the ER, Golgi, and vesicles exhibit a rhythm that peaks roughly around the night-to-day transition. Here we found that *Bmal1*^hep-/-^ liver secretes less albumin, and that *Bmal1*^hep-/-^mice and mice exposed to chronic jet lag exhibit a drop in total serum protein abundance during the inactive, fasting phase near ZT8. Albumin and a few other liver-secreted proteins account for most of the total protein content of the blood^1^. Therefore, timing of hepatic secretion may be critical to stabilize total serum protein abundance over the course of a day. For all proteins, we found a weak correlation between liver protein abundance and secretion according to time-of-day, but a moderate correlation according to disruption of *Bmal1*. Thus, the effect of clock disruption on the hepatic proteome can be observed in the secretome to a considerable degree. It is unlikely that one rhythm dictates the timing of all secreted proteins given the various control mechanisms that govern expression and secretion^18^. Measurements of protein trafficking through the different secretory pathways will be important to fully map the temporal aspects of hepatic secretory functions.

Evidence also supports a role for feeding rhythms as a key component of circadian control of secreted proteins. In humans, mistimed eating and sleeping induced by laboratory-simulated nightshifts disrupted daily patterns of a significant of portion many plasma proteins^15^. In addition, feeding and fasting are known to shape the plasma proteome^46^. Regarding liver, combined transcriptomic and proteomic analyses showed that archetypal secreted proteins with rhythmic protein abundance in the liver tend to have nonrhythmic mRNAs^7^, implicating post-transcriptional regulation known to occur in response to feeding and fasting. For this group of proteins, rhythms were maintained in night-fed *Cry1^-/-^*;*Cry2^-/-^*mice, indicating that a strong daily feeding rhythm can establish rhythms of secreted proteins within the liver. We find that in *Bmal1*^hep-/-^ mice, which have an intact feeding-fasting rhythm, most WT time-dependent secreted proteins lose time-dependence. However, secretion data also indicated *de novo* time-dependent secretion for many proteins in *Bmal1*^hep-/-^. The hepatocyte clock may buffer against feeding- and fasting- induced secretion for certain proteins, yet this notion needs to be rigorously tested. Ultimately, we surmise that the hepatocyte clock and feeding rhythms synergize to organize protein secretion over the circadian cycle, as is the case for other core liver circadian functions.

Clock transcription factors are known to regulate specific hepatokines central to metabolic homeostasis. The clock repressor REV-ERBα represses FGF21 via a mechanism involving HNF6^10^. Competing with REV-ERBα at RORE motifs, RORα activates *Fgf21*^47^ and *Fgb*^48^ transcription. Adropin exhibits circadian features and is responsive to REV-ERB and ROR agonists^49^. The PAR bZIP clock protein DBP transactivates *Rbp4*^9^. These clock-targeted hepatokines serve various functions throughout the body; RBP4 lowers whole-body insulin sensitivity^9^, FGF21 mediates metabolic responses to nutrient status as an endocrine hormone^50^, and another hepatokine, ANGPTL8, facilitates synchronization of the liver clock to feeding via autocrine signaling^3^. Previous work has also highlighted how clock control of secreted proteins is involved in etiology of disease phenotypes; aberrant induction of hepatokines (e.g., LBP, ITIH3, IGFBP1) that stimulate catabolic processes and muscle wasting in cancer cachexia models is due to loss of REV-ERBα-mediated transcriptional repression^13^. Here, we found that temporal secretion of endostatin involves CRY-mediated repression of the gene that encodes the full-length collagen protein COL18A. In mouse tendon, rhythmic properties of the secretory and endocytic pathways affect a pool of newly synthesized collagen which ultimately maintains collagen homeostasis across the circadian cycle^12^. Notably, the CRY agonist KL001 reduces the accumulation of collagen induced by circadian disruption. Whether this temporal collagen turnover mechanism extends to the liver and how it might affect collagen-derived soluble signaling peptides like endostatin is unclear, yet our results demonstrate temporal regulation of multiple extracellular matrix proteins and specifically link CRY to COL18A1 expression.

Endostatin was originally pursued as a cancer treatment due to its anti-angiogenic properties that can slow tumor growth and metastasis^38^. It currently has FDA-approved indications for neovascular-related cancers alongside anti-VEGF therapy and is safe for use in humans. More recently, potent metabolic effects of endostatin were discovered. Chronic administration of recombinant endostatin^39^ can counteract body weight gain and insulin resistance phenotypes of diet-induced obesity, in part, by inhibiting adipogenesis. It was shown that 3T3-L1 adipocytes internalize endostatin, which then interacts with the RNA-binding protein SAM68, ultimately lowering mTORC1 pathway activity^39^.

Here, we define effects of endostatin that are temporally restricted, contributing to rhythmic metabolic physiology. Endostatin lowered mitochondrial respiration in adipocytes, consistent with downregulation of mitochondrial genes in eWAT after injection of recombinant endostatin. Impaired adipocyte differentiation is shown to coincide with reduced mitochondrial respiration^45^, and respiratory rate may influence adverse metabolic outcomes, as adipocytes from individuals with obesity-related metabolic dysfunction exhibit higher mitochondrial respiration^51^. Additionally, endostatin altered the expression of lipid handling genes in eWAT and increased β-adrenergic-stimulated lipolysis while lowering lipid stores in adipocytes. Notably, endostatin did not affect insulin-stimulated glucose uptake. These results align with heightened endostatin secretion during the inactive, fasting phase and the temporally-restricted response of adipose tissue to the same window, indicating that endostatin acts not only to reduce adipogenic potential^39^ but also enhance lipolytic activity of adipocytes. Previous studies showed that hepatic COL18A1 protein displays diurnal rhythmicity, with a peak near ZT2 under both day-restricted and night-restricted feeding^52^, further substantiating our findings that COL18A1 and endostatin are regulated by the clock and aligned to the inactive, fasting phase. Whether post-translational modifications contribute another layer of temporal regulation to COL18A1 or endostatin remains to be seen. Endostatin may elicit these additional effects through known endostatin-targeted pathways such as mTOR^39^ or VEGF^53^ or yet-to-be-identified mechanisms. The underlying causes for the observed sex difference in response to endostatin remain to be determined.

Stromal vascular fraction progenitor cells that give rise to mature adipocytes exhibit a diurnal rhythm in adipogenesis that declines from ZT0 to ZT12 during the fasting phase^54^. Our secretion timing and circulating abundance data that show endostatin elevation over this time support a model in which endostatin contributes to this physiological rhythm as an inter-organ signaling factor. Notably, high-fat diet induces a constantly high rate of adipogenesis throughout the circadian cycle^54^, contributing to adverse lipid phenotypes. These observations raise the possibility that chronotherapeutic strategies may further enhance the anti-obesogenic effects of recombinant endostatin^39^ by aligning treatment time with adipocyte physiology^54^.

Ex vivo approaches are widely used to study liver function^55^ and are demonstrated to reflect the in vivo state of the liver^56^, including circadian phase^57^ and protein secretion^58^. In this study, the strengths of the acute ex vivo approach include circumventing phenotypic changes that arise during culture, preserving anatomical structures involved in secretion, retaining all hepatic cell types, capturing multiple mechanisms of secretion, and enabling rapid completion to maintain tissue integrity. This approach also has some inherent limitations. Constraints on the diffusion of oxygen and nutrients could induce hypoxia or stress leading to non-physiological secretion, yet measures of tissue oxygenation, metabolic rate, membrane integrity, and cell viability were all within reason. In addition, secreted proteomes were of similar composition to those generated from precision-cut liver slices, a preeminent approach to study protein secretion. Ultimately, we demonstrate the utility of the approach to identify mechanisms of temporal inter-organ crosstalk.

## METHODS

### Animals

Animal experiments were conducted in accordance with the National Research Council’s Guide for the Care and Use of Laboratory Animals and all experiments were conducted with approval from local Institutional Animal Care and Use Committee. ARRIVE (Animal Research: Reporting of In Vivo Experiments) guidelines were also followed wherever possible. Wild type C57BL/6J background mice ages 8 to 16 weeks were utilized for *ex vivo* secretion studies and isolation of primary hepatocytes throughout. Male and female mice were used for main experiments. Male mice were utilized for all other experiments, unless otherwise noted. Mice were group housed and fed *ad libitum* with vivarium chow. Unless otherwise indicated, livers were harvested at ∼ZT8 under a standard 12 h light: 12 h dark cycle. *Bmal1*-stop-FL mice have been described previously^59^. *Bmal1*-stop-FL mice were crossed with the Alfp-Cre line to generate mice with reconstitution of *Bmal1* in hepatocytes^41^. Experimental genotypes were: 1. wild type (WT) – *Bmal1^wt/wt^*, *Alfp-*cre^tg/0^; 2. *Bmal1* whole-body knockout – *Bmal1^stop-FL/stop-FL^*, *Alfp-*cre^0/0^; and 3. *Bmal1* hepatocyte-reconstituted mice - *Bmal1^stop-FL/stop-FL^*; *Alfp*-cre^tg/0^. *Bmal1* hepatocyte-specific knockout mice (*Bmal1*^hep-/-^) were generated by crossing *Bmal1*-flox mice^60^ with Alfp-Cre. Experimental genotypes were: 1. wild type (WT) – *Bmal1^flox/flox^*, *Alfp-*cre^0/0^; and 2. *Bmal1*^hep-/-^– *Bmal1^flox/flox^*, *Alfp-*cre^tg/0^. For chronic jet lag and endostatin injection studies, mice C57BL/6J mice (strain #000664) were purchased from The Jackson Laboratory. Mice were allowed at least one week adaptation upon arrival. For the fasting-refeeding experiment, food was removed from the cages at ZT12, checking for any pellet pieces in the bedding. Mice then proceeded to the ex vivo secretion protocol at ZT24/0 (fasting group) or were readministered food ad libitum from ZT24/0 to ZT4 and then proceeded to the ex vivo secretion protocol at ZT4 (refed).

### Protein secretion from the liver *ex vivo*

Mice were anesthetized with 2.5% isoflurane. The abdominal cavity was exposed, the hepatic portal vein was clipped with scissors, and then the inferior vena cava was immediately cannulated using a butterfly needle. The liver was then perfused with PBS at 2 mL/min for 2 minutes for a total volume of 4 mL. After perfusion, a ∼100 mg piece of the left liver lobe was removed and placed in a 50 mL conical tube containing 5 mL of CD CHO media (Gibco, 10743029) pre-bubbled with carbogen to ensure oxygen availability during tissue transfer from the animal facility to the laboratory, which occurred within ∼3-5 min. The liver was then incubated in a water bath for 45 min at 37°C while the media was continuously bubbled with carbogen. Following incubation, the liver was frozen at - 80 for downstream applications and the media was centrifuged at 2,000 x g for 5 min at 4°C to pellet cells/cellular debris. The supernatant was then sonicated using the Bioruptor Pico (Diagenode) 6 times for 20 sec on and 30 sec off on the 15 mL setting. The secreted fraction was then concentrated in the Amico Ultra 15 Centrifugal Filters (Millipore) at 4°C for 45 minutes at 4,000g to ∼20-fold concentration. Total protein was measured by BCA prior to downstream applications. For the brefeldin A experiments, input for western blot was normalized by starting liver weight.

### Western blot

#### Sample preparation

Livers were homogenized on ice in RIPA buffer (Thermo Scientific, J62524.AE) supplemented with a protease inhibitor cocktail (Thermo Scientific, 78429). The samples were incubated on ice for 10 min to lyse cells and then sonicated with 4 cycles of 20 sec on, 30 sec off on 1.5 mL setting using the Bioruptor Pico (Diagenode). The samples were then centrifuged at max speed at 4°C for 10 min and the supernatant was collected. Protein concentration was determined using the Pierce BCA Protein Assay (Thermo Scientific, 23225). Concentrated protein samples from the secreted fraction entered the protocol at this step. Proteins from conditioned media of primary hepatocytes were prepared as follows: conditioned media from three 100 mm dishes was pooled and centrifuged at 2,000 x g for 5 min at 4°C to pellet cells/cellular debris, sonicated as in the ex vivo method, and then the supernatant was concentrated with 10 kDa centrifugal filters (Millipore, 0000338873) to final volume of 250 µL. To enhance detection of potentially lowly abundant secreted proteins, concentrated protein samples were treated with the Proteospin Abundant Serum Protein Depletion Kit (Norgen Biotek, 17300), then concentrated down further with amicon ultra 0.5 mL centrifugal filter tubes (Millipore, UFC500396) and finally, protein concentration was determined as described above. All western samples were either loaded by protein mass, or by liver mass for the brefeldin a experiments. The appropriate volume of 6X loading buffer (VWR, J61337) was added and PBS was used as a diluent to reach the appropriate volume per well.

#### Blotting

Protein was separated on 4-20% gradient gels by SDS-PAGE and subsequently transferred to a PVDF membrane using the Trans-Blot Turbo Transfer System (Bio-Rad, 1704150). The amount of starting protein ranged from 15 ug to 30 ug depending upon the application. Blots were blocked with 5% milk in TBS with 0.1% tween-20 (TBST) at room temperature for 1 hour. Primary antibodies were diluted in 5% milk TBST and exposed to blots by overnight incubation at 4°C: Albumin (Bethyl Laboratories, A90-134A); H3 (Cell Signaling, 14269); SOD2 (Proteintech, 24127-1-AP); CES2 (Novus Biologicals, NBP1-91620); Normal Mouse IgG (Cell Signaling, 68860); FGB (ProteinTech, 16747-1-AP; BMAL1 (Abcam, ab93806); Col18a1/endostatin (Santa Cruz, sc-32720); TAT (Santa Cruz, sc-376292); APOC4 (Santa Cruz, sc-134263); HIF-1α (Cell Signaling, 36169); β-Actin (Abcam, ab8226). Blots were washed 3x for 5 min with TBST and then incubated with secondary antibodies conjugated to HRP (Millipore Sigma, AP160P and 12-348) for 1 hour at room temperature. Blots were again washed 3x for 5 min with TBST and then incubate with HRP substrate (Millipore, WBLUC0500) for 5 min at room temperature, followed by visualization using the ChemiDoc Imaging System (Bio-Rad). Blots were quantified using ImageJ software and normalized to total protein using the stain-free gel feature of Mini-Portein TGX gels (Bio-Rad, e.g., 4568095). HIF-1α/2 α Control Cell Extracts (Cell Signaling, 94790) were used as controls for HIF-1 α blots. Uncropped blots are presented in the Source Data File or Supplementary Information File.

### Cell culture

#### AML12 cells

AML12 cells, purchased from ATCC (CRL-2254), were cultured in DMEM:F12 (ATCC, 30-2006) supplemented with 10% FBS (Corning, 35-015-CV), 1% penicillin-streptomycin, insulin-transferrin-selenium supplement (Corning, 25-800-CR), and dexamethasone (40 ng/mL). Cells were plated at confluency of 40% in 96-well plate and the following day were transfected with the respective plasmids (*Bmal1, Clock, Cry1, Cry2*, *fgb-luc, col18a1-luc, and scarmble-luc*) at their corresponding doses according to the Lipofectamine 3000 Transfection Kit (Invitrogen, L3000001) with the exception of incubating the cells in OPTI-MEM for 4 hours after adding transfection reagents. Following the replacement of media, cells were transfected for 24 hours before measuring luciferase activity via the Dual-luciferase Reporter 1000 Assay System (Promega) according to the manufacturer’s protocol.

#### Primary hepatocytes

Primary hepatocytes were isolated from male WT or *Bmal1*^hep-/-^ mice, ages 8 to 16 weeks, using the Liver Perfusion Kit, mouse and rat (Miltenyi Biotec, 130-128-030) and the gentleMACS Octo Dissociator with Heaters (Miltenyi Biotec, 130-096-427) according to the manufacturer’s instructions. Primary hepatocytes were plated at a density of 1 X 10^6^ cells/100 mm dish in DMEM (Corning, 10-013-CV) supplemented with 5% FBS and 1% Penicillin/Streptomycin. For experiments featuring unsynchronized cells, the following day, for conditioned media collection, the media was replaced with serum-free media for 4 h before collecting and concentrating as described in the Western blot section. For synchronization experiments, the following day, cells were synchronized by 1 h exposure to 100 nM dexamethasone. Starting 12 h post-synchronization, secreted proteins were collected in 4 h windows from separate wells (i.e. not the same well repeatedly) out to 36 h (according to serum-free media exposure as described above). At the time of conditioned media collection, cells were also collected for qPCR measurements.

### 3T3-L1 adipocytes

3T3-L1 pre-adipocytes were cultured and differentiated as previously described^61^ with slight modifications. 3T3-L1 pre-adipocytes were expanded in DMEM, high-glucose (Corning, 10-013-CV), with 10% Fetal Calf Serum (FCS) and 1% P/S. Cells were plated and allowed to remain at confluence for 48 h before inducing differentiation. For inducing differentiation, FCS was changed to Fetal Bovine Serum (FBS), and IBMX (50 mM), Insulin (8 µg/mL), and Dexamethasone (1 µM) were added. The media was then not changed for 4 days. After day 4, the induction media was changed to an insulin media which contained DMEM, high-glucose, 10% FBS, 1% P/S, and 8 µg/mL Insulin. Experiments were performed on day 10 following the start of differentiation.

### Luciferase experiments

AML12 cells were transfected with plasmid DNA at 75% confluency using Lipofectamine 3000 Transfection Reagent (Invitrogen, #L3000001). Luminescence was measured 48 hours later using the Varioskan LUX Multimode Microplate Reader (Thermo Fisher Scientific) and Dual-Glo Luciferase Assay System (Promega, E2940), wherein firefly luciferase signal from the experimental luciferase reporter plasmid was normalized to the control renilla luciferase signal.

### Plasmids

Luciferase reporter constructs were synthesized by inserting promoter sequences into pGL3-basic vectors at the Xhol (5’ restriction enzyme) and HindIII (3’ restriction enzyme) sites. The Eukaryotic Promoter Database^62,63^ was used to retrieve the sequence from -2 kb to +1 kb relative to the transcription start site (TSS): *Col18a1* = >FP014233 Col18a1_1:+U EU:NC; range -2000 to 1000; *Fgb* = >FP003937 Fgb_1 :+U EU:NC; range -2000 to 1000. The scrambled control sequence was generated from the *Fgb* promoter sequencing using the Sequence Manipulation Suite Shuffle DNA tool. For the *Fgb* E-box mutant construct, putative E-box elements were identified using the Eukaryotic Promoter Database Search Motif Tool (Library: JASPAR core 2018 vertebrates; Motif: Arntl; From:-2000; To:1000 bp relative to TSS and a cut-off [p-value] of: 0.001). Full plasmid sequences, maps, and quality control information are available in Supplementary Data 4. These plasmids are available upon request. All other plasmids were purchased from Addgene: Cry1 - #110298; Cry2 - #31283; Bmal1 - #31367; Clock - #31366.

### Seahorse assay

Seahorse assay was conducted as we have done so previously^8^ with slight modifications to the manufacturer’s protocol for XF Mito Stress Test Kit (Agilent, 103015-100). 3T3-L1 adipocytes were plated at a density of 10,000 cells per well in seahorse cell culture plates and differentiated as described above. 1 h prior to the start of the assay, insulin media was removed and cells were incubated in Seahorse XF Base Medium (Agilent) with 10 mM glucose, 1mM pyruvate, and 2 mM L-glutamine added, pH-adjusted to 7.4. The standard mito-stress test was performed, with OCR and ECAR measurements at three time points before injections of oligomycin (1 µM), FCCP (2 X 1 µM), and rotenone (0.5 µM) + antimycin A (0.5 µM). Injections were followed by 3 minutes of mixing and a 3-minute measurement period, which was repeated 3 times for each stage. Each well was normalized by total protein as measured by Pierce BCA Assay (ThermoScientific, 23223). Basal respiration was determined by subtracting the average of the rotenone/antimycin A readings from the Basal OCR average. Maximal respiration, similarly was determined by subtracting the average measurements after the second FCCP injection from the rotenone/antimycin average. Glycolytic compensation was calculated as post-oligomycin ECAR minus baseline ECAR. Stressed glycolysis represents the 2^nd^ FCCP ECAR minus glycolytic compensation. PBS was used as control treatment.

### BODIPY measurement

Cellular neutral lipid content was determined using BODIPY 493/503 (TargetMol, T36957). Cells were fixed in 4% PFA for 20 minutes at room temperature and rinsed 3 times with PBS. BODIPY and Hoechst (ThermoScientific, 62249) were incubated for 10 minutes at room temperature at a concentration of 0.1 mg/mL and 2 mM, respectively. Cells were washed three times with PBS and then fluorescence readings at 493/511 and 350/461 were recorded for BODIPY and Hoechst, respectively, using the Varioskan LUX Multimode Microplate Reader (ThermoScientific) BODIPY was normalized to cell number by Hoechst intensity. 5 ug/mL BSA was used as control treatment.

### Lipolysis assay

Lipolysis was measured using the 3T3-L1 Lipolysis Kit (zenbio, LIP-3-NCL1) according to the manufacturer’s protocol. 3T3-L1 adipocytes were cultured and differentiated as described in Cell culture. The day of the experiment, insulin was washed out for 2 h using maintenance media (DMEM, high glucose, 10% FBS, and 1% P/S). Following the washout, cells were treated for 2 h with forskolin (5 uM) or .01% DMSO vehicle, and/or 5 ug/mL Endostatin or PBS control. After 2 h, non-esterified fatty acid concentration was measured in the media.

### Glucose uptake assay

Glucose uptake was measuring using the Glucose Uptake-Glo Assay (Promega, J1342) according to the manufacturer’s protocol. 3T3-L1 adipocytes were cultured and differentiated as described in Cell culture. The day of the experiment, insulin was washed out for 2 h using maintenance media (DMEM, high glucose, 10% FBS, and 1% P/S). Following the washout, cells were treated with 5 µg/mL endostatin or PBS control for 2 h, prior to 100 nM insulin incubation for 1 h. Endostatin remained in the media during insulin treatment. After 1-hr insulin incubation, glucose uptake was measured via luminescence readout using the Varioskan LUX Multimode Microplate Reader (Thermo Fisher Scientific).

### Cathepsin L activity

Cathepsin L activity was measured using the Innozyme Cathepsin L Activity Kit, Fluorogenic (Millipore Sigma, CBA023-1KIT) according to the manufacturer’s protocol, starting from 50 ug of liver protein extracts prepared as in Proteomics (dataset 2).

### Endostatin ELISA

Circulating endostatin was quantified using the Mouse Endostatin ELISA Kit (Abcam, ab263894) according to the manufacturer’s instructions. Serum was prepared by placing blood on ice for at least 10 min, followed by centrifugation for 15 min at 3,000 x g at 4°C. Plasma was prepared as follows: blood was collected, placed on ice in a Microvette CB 300 K2E collection tube (Starstedt, 16.444.100), and centrifuged at 1,500 x g for 15 min at 4°C. The supernatant was then aliquoted and stored at -80 for downstream assays.

### Hematoxylin & eosin staining

Formalin-fixed, paraffin-embedded slides were deparaffinized in serial washes of xylene (3x; 5 mins.), 100% ethanol (2x; 1 min.), 95% ethanol (1x; 1 min.) and deionized water (1x; 1 min.). Hematoxylin & eosin staining was performed on a Leica Autostainer XL where slides were sequentially incubated in modified Harris hematoxylin (filtered; 1 min.), acid alcohol (20 sec.), Scott’s tap water solution (15 sec.), 95% ethanol (1 min.), and eosin y alcoholic solution (2 min.), with tap water washing between each step (except ethanol and eosin). Subsequently, slide were dehydrated in ethanol and xylene series, and coverslipped using Cytoseal XYL permanent mounting media. 10 images were taken per biological replicate at 40X, and the “point” tool and the “CellCounter” plugin in imageJ were used to count the number of swollen nuclei/cytoplasm per field as well as the number of total cells, respectively, after blinding the researcher to treatment. The percentage of swollen nuclei/cytoplasm was calculated by dividing by the total number of nucleated cells per image.

### Proteomics (dataset 1)

#### Data acquisition

Total protein from each sample was reduced, alkylated, and purified by chloroform/methanol extraction prior to digestion with sequencing grade modified porcine trypsin (Promega). Tryptic peptides were then separated by reverse phase XSelect CSH C18 2.5 um resin (Waters) on an in-line 150 x 0.075 mm column using an UltiMate 3000 RSLCnano system (Thermo). Peptides were eluted using a 60 min gradient from 98:2 to 65:35 buffer A:B ratio (Buffer A = 0.1% formic acid, 0.5% acetonitrile; Buffer B = 0.1% formic acid, 99.9% acetonitrile). Eluted peptides were ionized by electrospray (2.2 kV) followed by mass spectrometric analysis on an Orbitrap Eclipse Tribrid mass spectrometer (Thermo). To assemble a chromatogram library, six gas-phase fractions were acquired on the Orbitrap Eclipse with 4 m/z DIA spectra (4 m/z precursor isolation windows at 30,000 resolution, normalized AGC target 100%, maximum inject time 66 ms) using a staggered window pattern from narrow mass ranges using optimized window placements. Precursor spectra were acquired after each DIA duty cycle, spanning the m/z range of the gas-phase fraction (i.e. 496-602 m/z, 60,000 resolution, normalized AGC target 100%, maximum injection time 50 ms). For wide-window acquisitions, the Orbitrap Eclipse was configured to acquire a precursor scan (385-1015 m/z, 60,000 resolution, normalized AGC target 100%, maximum injection time 50 ms) followed by 50x 12 m/z DIA spectra (12 m/z precursor isolation windows at 15,000 resolution, normalized AGC target 100%, maximum injection time 33 ms) using a staggered window pattern with optimized window placements. Precursor spectra were acquired after each DIA duty cycle.

#### Data analysis

Following data acquisition, data were searched using Spectronaut (Biognosys version 18.5) against the UniProt *Mus musculus* database (April 2023) using the directDIA method with an identification precursor and protein q-value cutoff of 1%, generate decoys set to true, the protein inference workflow set to Quant 2.0, inference algorithm set to IDPicker, quantity level set to MS2, cross-run normalization set to false, and the protein grouping quantification set to median peptide and precursor quantity. Protein MS2 intensity values were assessed for quality using ProteiNorm^64^. The data was normalized using VSN and analyzed using proteoDA (DOI: 10.21105/joss.05184) to perform statistical analysis using Linear Models for Microarray Data (limma)^65^. A filtering protocol removed proteins from the dataset if 2/3 replicates had missing values.

### Proteomics (dataset 2, WT and *Bmal1*^hep-/-^)

#### Data acquisition

Liver and secretome samples were separately blocked and randomized for sample preparation and mass spectrometry analysis. Liver homogenates were vortexed in buffer containing 5% SDS/50 mM triethylammonium bicarbonate (TEAB) in the presence of protease and phosphatase inhibitors (Halt; Thermo Fisher Scientific) and nuclease (Pierce™ Universal Nuclease for Cell Lysis; Thermo Scientific). Aliquots corresponding to 100 µg protein (EZQ™ Protein Quantitation Kit; Thermo Scientific) were reduced with tris(2-carboxyethyl)phosphine hydrochloride (TCEP); alkylated in the dark with iodoacetamide and applied to S-Traps (mini; Protifi) for tryptic digestion (sequencing grade; Promega) in 50 mM triethylammonium bicarbonate (TEAB). Peptides were eluted from the S-Traps with 0.2% formic acid in 50% aqueous acetonitrile, taken to dryness in a SpeedVac (Thermo Fisher Scientific) vacuum concentrator, reconstituted in starting HPLC mobile phase (3% B, see below) and quantified using Pierce™ Quantitative Fluorometric Peptide Assay (Thermo Fisher Scientific); samples were diluted as needed to achieve a concentration of 0.4 µg/5 µl. For the secretome samples, 50-µl aliquots were mixed with 50 µl of 10% SDS/50 mM TEAB in the presence of protease and phosphatase inhibitors and subsequently reduced/alkylated and digested with trypsin as described above except that micro S-Traps were used. Dried peptides were taken up in 20 µl of starting HPLC mobile phase. Digests were analyzed by capillary HPLC-electrospray ionization data-independent acquisition mass spectrometry (DIA-MS) on a Thermo Scientific Orbitrap Exploris 480. On-line separation was accomplished with a Vanquish Neo UHPLC system (Thermo Scientific): column, PepSep (Bruker; ReproSil C18, 15 cm x 150 µm, 1.9 µm beads); mobile phase A, 0.5% acetic acid (HAc)/0.005% trifluoroacetic acid (TFA) in water; mobile phase B, 90% acetonitrile/0.5% HAc/0.005% TFA/9.5% water; gradient 3 to 42% B in 60 min; flow rate, 0.4 μl/min. Separate pools were made of the digests of the 32 liver and secretome samples: secretome, equal volumes; cells, equal quantities of peptides. Aliquots of the pools (secretome, 5 µl; cells, 2 µg/5 µl) were analyzed using three stages of gas-phase fractionation (400–600 m/z; 600–800 m/z; 800–1000 m/z) and 4-m/z windows (30k resolution for precursor and product ion scans).

### Data analysis

The resulting three data files were used to create an empirically-corrected DIA chromatogram library by searching against a Prosit-generated predicted spectral library^66^ based on the UniProt mouse reference database [UniProt_Mouse_ref 10090_20220216 (21,986 sequences; 11,739,283 residues)]. Injections of the digests of the individual experimental samples were: secretome, 5 µl of digest; cells, 2 µg peptides/5 µl. HPLC conditions were the same as described above for generation of the chromatogram library. MS data for the experimental samples were acquired using 8-m/z windows (400–1000 m/z; staggered; 30k resolution for precursor and product ion scans) and searched against the chromatogram library. Scaffold DIA (v4.1.0; Proteome Software) was used for all DIA-data processing: fixed modification, cysteine carbamidomethylation; proteolytic enzyme, trypsin with one missed cleavage allowed; peptide mass tolerance, ±10.0 ppm; fragment mass tolerance, ±10.0 ppm; charge states, 2+ and 3+; peptide length, 6–30. Peptides identified in each sample were filtered by Percolator ^67^ to achieve a maximum FDR of 1%. Individual search results for cells and secretome were separately combined and peptide identifications were assigned posterior error probabilities and filtered to an FDR threshold of 1% by Percolator. Peptide quantification was performed by Encyclopedia ^68^ based on the three to five highest quality fragment ions. Only peptides that were exclusively assigned to a protein were used for relative quantification. Two samples from the secretome dataset displayed a clear divergence in Retention Time (RT) deviation as compared to the other samples, therefore, they were removed from the data analysis.

### RNA sequencing

Total RNA was extracted from 100 mg of eWAT which was homogenized with a mortar and pestle in liquid nitrogen and separated with 1 mL QIAzol Lysis Reagent (Qiagen, 79306) with repeated vortex steps to dissociate lipid phase. The lipid layer was removed from the surface after 5 minute incubation at room temperature and a 12,000 x g spin for 10 minutes at 4°C. The clear homogenate was then transferred to a new tube, and 200 µL of chloroform was added. Samples were vortexed vigorously, incubated for 10 minutes at room temperature and centrifuged for 15 minutes for 12,000 x g at 4°C. RNA was then collected from the aqueous phase and 500 µL isopropanol was added. RNA was vortexed, incubated for 10 minutes, and then centrifuged as per the first centrifugation step. Following centrifugation, the supernatant was removed and the pellet was washed with 75% ethanol with a final centrifugation at 7500 x g for 5 minutes at 4°C. Once RNA was collected, cleanup was performed using the RNeasy Plus Mini Kit (Qiagen, 74134) according to manufacturer’s recommendations.

The quality of Total RNAs was checked by Agilent Tape Station (Agilent Technologies, Santa Clara, CA), and RNAs passed QC with RIN >7. Approximately 500ng Total RNA was used for Total RNA-seq library preparation by following the NEB Directional RNA library preparation guide with NEBNext rRNA depletion kit v2 (human/mouse/rat) (New England Biolabs, Ipswich, MA). The first step in the workflow involved the depletion of rRNA by hybridization of complementary DNA oligonucleotides, followed by treatment with RNase H and DNase to remove rRNA duplexed to DNA and original DNA oligonucleotides, respectively. Following rRNA removal, the left RNA was fragmented into small pieces using divalent cautions under elevated temperature and magnesium. The cleaved RNA fragments were copied into first strand cDNA using reverse transcriptase and random primers. This was followed by second strand cDNA synthesis using DNA Polymerase I and RNase H. Strand specificity was achieved by replacing dTTP with dUTP in the Second Strand Marking Mix (SMM). The incorporation of dUTP in second strand synthesis effectively quenches the second strand during amplification, since the polymerase used in the assay will not incorporate past this nucleotide. These cDNA fragments then went through an end repair process, the addition of a single ‘A’ base, and then ligation of the adapters. The products were then purified and enriched with PCR to create the final RNA-Seq library. After RNA-seq libraries were subjected to quantification process, pooled for subsequent 150bp paired end sequencing run with Illumina NovaSeq platform. After the sequencing run, demultiplexing was employed to generate the fastq file, with the average of ∼42M reads per sample.

Salmon indexing (V.1.10.2) was performed using GenCode’s GRCm39 Ensemble Release 114 reference genome utilizing Ensembl GTF files. Quantification was corrected for GC bias using the –gcbias option. Quantification was performed with the – validateMappings option to ensure alignment accuracy. Transcript abundance estimates were imported using tximports (v.1.30.0), and tx2gene mapped the gene level abundance by Ensembl annotations. Gene counts were normalized an analyzed for differential expression using DESeq2 (v1.42.1). An adjusted p-value of < 0.05 (Benjamini-Hochberg FDR) and an absolute log2 fold change of 0.58 or -0.58 (50% change) were used as cutoffs for differential expression.

### Gene ontology analysis

Gene ontology analysis was performed using the Database for Annotation, Visualization and Integrated Discovery (DAVID). GOTERM_DIRECT charts were utilized. Complete functional annotation charts can be found in Supplementary Data 2.

### Quantitative PCR

RNA was extracted using the RNeasy Plus Mini Kit (Qiagen, 74136) and cDNA was prepared from 1 ug RNA using the Maxima First Strand cDNA Synthesis Kit (Thermo Scientific, K1641). Quantitative real-time PCR was performed on a QuantStudio 5 with PowerUp SYBR Green Master Mix (Applied Biosystems, A25742) and normalized to *18s* control. Primer sequences were identified using the Harvard PrimerBank Database^69^ and are listed in Supplementary Data 5.

### Chronic jet lag

Chronic jet lag was performed as we have done so previously^70^. Please see this reference for the precise tissue collection days. Briefly, male mice were acclimated to the animal facility for at least 1 week and group housed in a standard home cage environment inside Circadian Cabinets (Actimetrics) at room temperature (23.9°C±1.5) with humidity monitoring (∼30%) and light intensity at ∼100 lux at cage level. Light schedules were fully automated and mice were only disturbed for husbandry activities, e.g. bedding changes, every 2 weeks. Between 4 and 6 weeks-of-age, mice were randomly assigned to one of two paradigms, which were impacted for a ∼3 month period: 1) control 12 hour light: 12 hour dark (LD); 2) 8-hour advance twice per week for 3 months (8 h adv 2x/wk-CJL). Locomotor activity was recorded with the imbedded cage infrared sensors (z-y ambulatory counts). Data from a 2-week period at the end of the paradigm was analyzed with ClockLab V.6 software (Actimetrics). Calculated parameters were generated using the following analysis features: 1) Activity Profile (Least Squares sine-wave fit, F statistics and p-value estimation [Cosinor]), 2) Periodogram (Chi-squared, F, and Lomb-Scargle), and 3) Automated detection of activity onsets for period estimation.

### Recombinant endostatin injection

15-week-old male and female C57BL/6J mice were either housed in circadian cabinets (Actimetrics) for two weeks in normal light-dark (lights on at 7 AM) or inverted light-dark (lights on at 7PM) prior to ZT6 and ZT18 injections respectively. Animals were injected via the tail vein with either 100 µg of recombinant human endostatin (PeproTech, Cat#150-01) or PBS vehicle control and food was removed for 2 hours prior to tissue collection. Blood glucose was measured with the nova max plus glucose and ketone monitor (Nova, B220714192XP) and glucose test strips (Nova, 43523).

### Statistical analyses

All data are mean ± S.E.M. unless otherwise indicated. For each experiment, sample size, statistical test, and significance threshold can be found in the figure legend and/or main text. Statistical analyses of large-scale datasets are described within the corresponding methods section. Field standards were used to determine appropriate sample sizes for proteomic analyses and circadian experiments. In instances where sample size is not explicitly stated, experiments were repeated at least three times, and one representative replicate is shown. Data were analyzed in Prism 11.0.0 (93) software (GraphPad), ClockLab 6, ImageJ 1.53t, or R Studio 2026.01.0+392 unless otherwise noted. Statistical assumptions were assessed for the appropriateness of ANOVA. Kolmogorov-Smirnov tests were used to assess Gaussian distribution (normality). Bartlett’s tests were used to assess homogeneity of variance. Observations within each group were independent. Mice were randomly assigned groups and sampled.

### Data availability

The proteomics data generated in this study have been deposited to the ProteomeXchange Consortium using the Proteomics Identifications Database (PRIDE) under accession codes PXD063542 (http://proteomecentral.proteomexchange.org/cgi/GetDataset?ID=PXD063542) and PXD066772 (http://proteomecentral.proteomexchange.org/cgi/GetDataset?ID=PXD066772). The RNA-sequencing data have been deposited to Gene Expression Omnibus (GEO) under accession code GSE305072 (https://www.ncbi.nlm.nih.gov/geo/query/acc.cgi?acc=GSE305072). Source data are provided with this paper in the Source Data File and Supplementary Information.

## ACKNOWLEDGEMENTS

For proteomics support, we thank the IDeA National Resource for Quantitative Proteomics and the San Antonio Institutional Mass Spectrometry Laboratory (UTHSCSA), with expert technical assistance of Sammy Pardo and Dana Molleur, under the direction of Susan T. Weintraub, Ph.D, and Min Kyu Kim, Ph.D.. For sequencing support, we thank the Genome Sequencing Facility of the Greehey Children’s Cancer Research Institute (UTHSCSA).

## FUNDING STATEMENT

The Koronowski lab is supported by the NIGMS under award number R35GM150618, the Max and Minnie Tomerlin Voelcker Fund, and the American Heart Association (25CDA1451928). Christopher Litwin is supported by the Translational Science Training (TST) T32 Program under award number T32TR004545 and by the NIDDK under award number F31DK143733. Kristi Dietert is supported by the NIA under award number F32AG096998. The IDeA National Resource for Quantitative Proteomics is supported under award number R24GM137786. The San Antonio Institutional Mass Spectrometry Laboratory is supported in part by NIH grant S10 OD030371-01A1 (S.T. Weintraub, for Orbitrap Exploris 480). The Genome Sequencing Facility is supported by NIH-NCI P30 CA054174, the NIH Shared Instrument grant S10OD030311 (S10 grant to NovaSeq 6000 System), and CPRIT Core Facility Award (RP220662). The lab of Lily Dong is supported in part by the NIH (R01DK134637) and the Baptist Health Foundation of San Antonio (2020 Strategic to Mission).

## AUTHOR CONTRIBUTIONS

C.L., L.Q.D., K.F.B., and K.B.K., conceived and designed the study. C.L. and K.B.K. wrote and edited the manuscript. C.L., Q.Z., I.T., Z.L., S.P., K.D., M.K., T.S., and J.R. performed experiments. S.H. performed bioinformatic analyses.

## COMPETING INTERESTS DECLARATION

The author declare no competing interests.

## Supplementary Information

### Supplementary Figures

**Supplementary Fig. 1.**
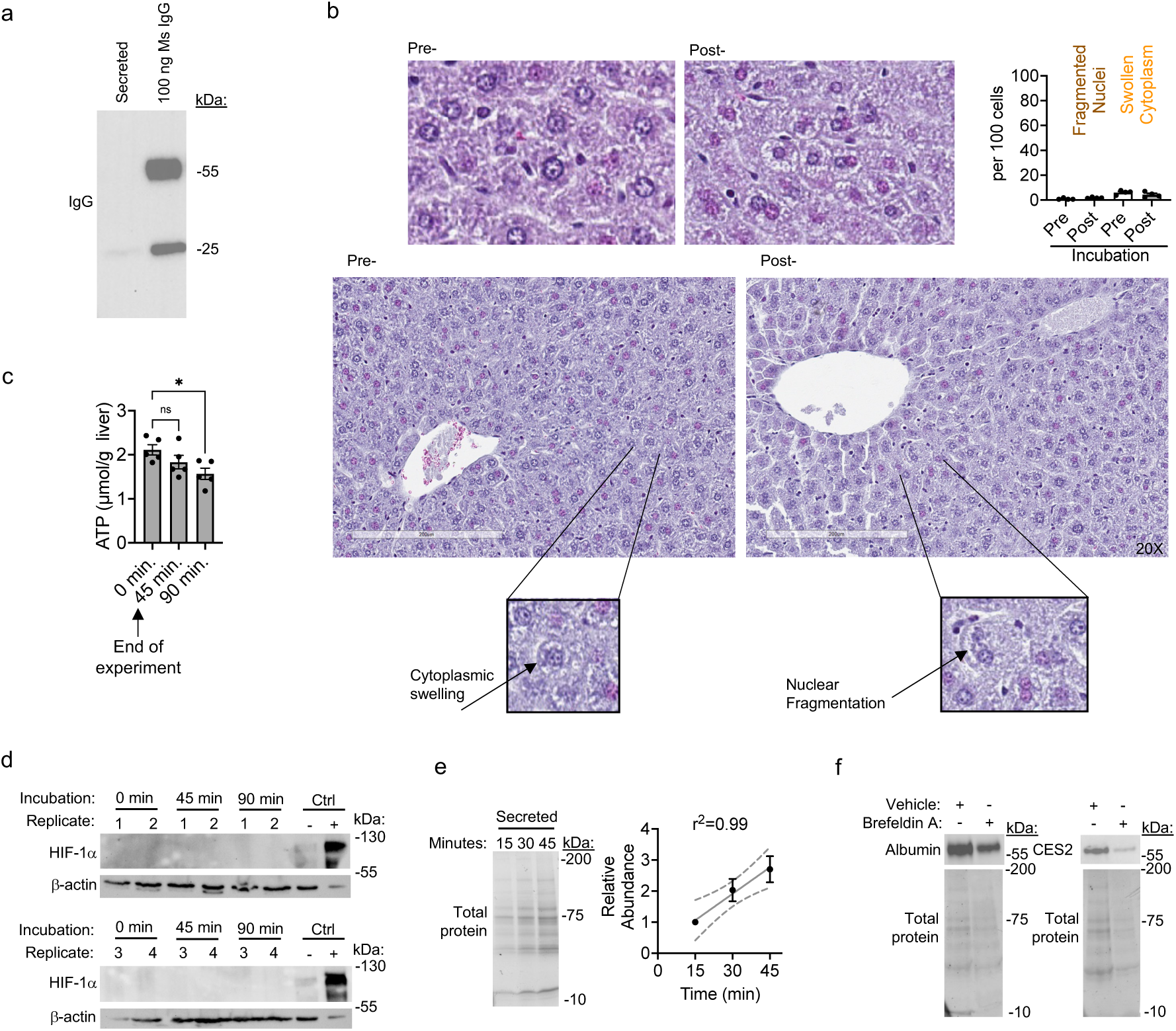
Data Related to Fig. 1. Data from ex vivo liver secretion experiments. (a) Representative Western blot showing minimal blood contamination in the secreted fraction. (b) Examples of H&E staining of pre- and post-incubation livers. N=4 quantification is shown in upper right corner. (c) ATP concentration in livers at the indicated incubation times. Note that all experiments in this study end at 45 min incubation, One-way ANOVA, Dunnett’s post hoc test, *p=0.0265, n=5. (d) Western blot from livers at the indicated incubation times. 80 ug protein loaded. + and - Ctrls = HIF-1a control extracts (Cell Signaling, #94790), n=4. (e) Left - Western blot of the secreted fraction from an incubation time course. Right – quantification of left, n=3. Pearson correlation r^2^ value shown. (f) Western blot examples for main Fig. 1c, wherein liver was incubated ex vivo with 2 mM Brefeldin A or 2% DMSO vehicle control for 45 min. CES2 – carboxylesterase 2. Source data for (a)-(f) are provided as Supplementary Source Data.

**Supplementary Fig. 2.**
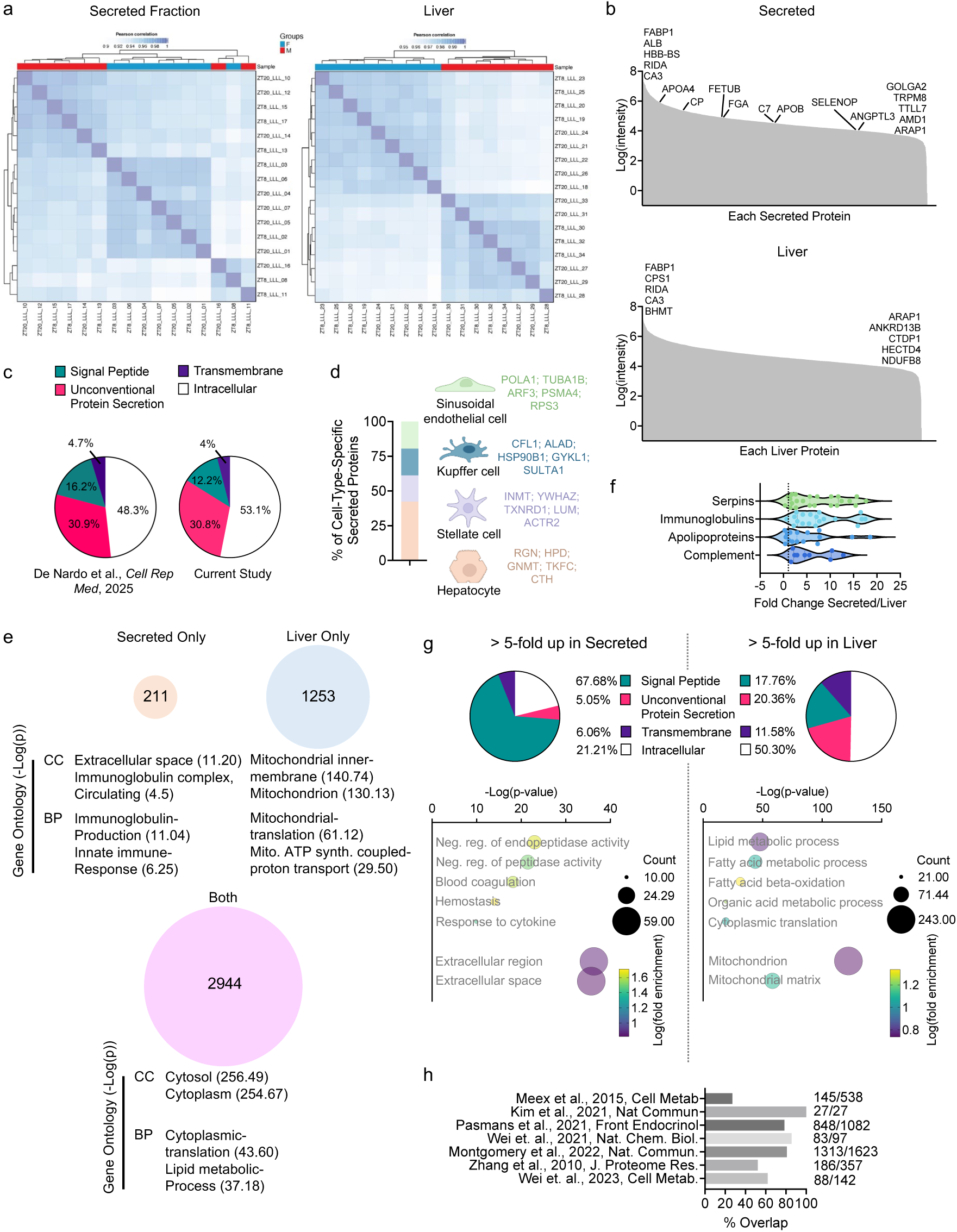
Data Related to Fig. 1. (a) Heatmap of Pearson correlation coefficients among samples showing an overall consistency among biological replicates. (b) Raw intensities of proteins quantified in the secreted and liver fractions showing the large dynamic range of quantification. The top 5 most and least abundant proteins are listed at the far left and far right, respectively. Notable secreted proteins are highlighted in between. (c) OutCyte classifications for secreted proteins from the current study and a reference study. (d) Specific analysis of cell-type-specific proteins. The top 5 most highly abundant proteins are listed to the right. Created in BioRender. Koronowski, K. (2026) https://BioRender.com/2tc2t15. (e) Overlap of proteins quantified in the liver and secreted fraction. Gene Ontology enrichment (DAVID) analysis of each group of proteins. Top 2 terms from Cellular Compartment (CC) and Biological Process (BP) are shown, with -log10 p-value in parentheses. (f-g) Fold change analyses. (f) classical plasma protein families showing enrichment in the secreted fraction compared to liver. (g) OutCyte classifications for proteins >5-fold enriched in either fraction. Bubble plots of Gene Ontology enrichment from DAVID, with top 5 terms from Biological Process and top 2 from Cellular Compartment. (h) Comparison of ex vivo secreted proteins from this study with the indicated published datasets. Values are (# of overlapping proteins) / (total from published dataset).

**Supplementary Fig. 3.**
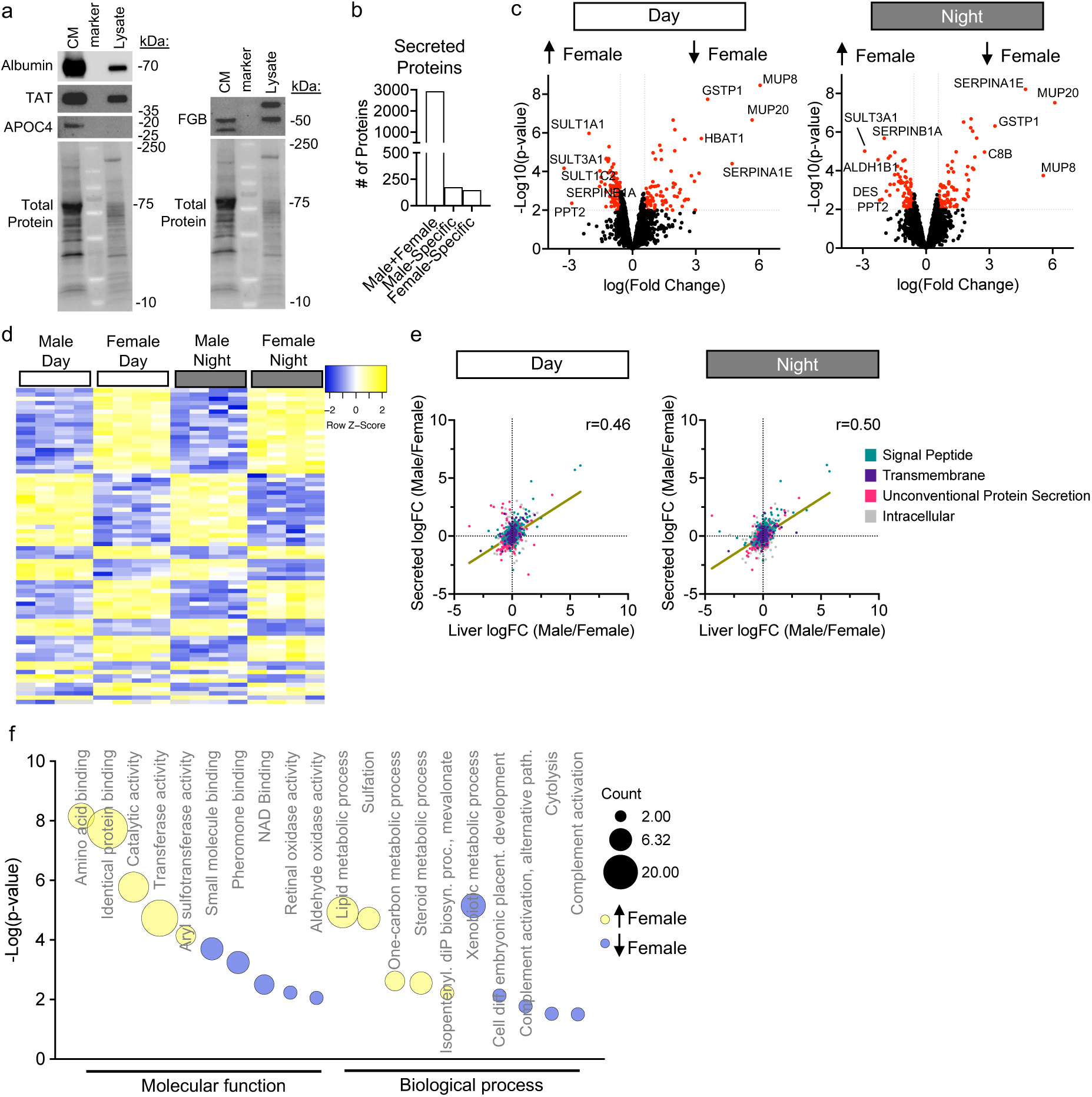
Data Related to Fig. 2. (a) Western blot from primary hepatocyte cultures. CM - conditioned media, 4 h incubation. TAT – tyrosine aminotransferase. APOC4 – apolipoprotein C-IV. FGB – fibrinogen beta chain. (b-f) Analysis of secreted fraction proteomes by sex. (b) # of sex-dependent and sex-independent secreted proteins. (c) Volcano plots showing sex-dependent secreted proteins. Red data points = Student’s t-test, p<0.01, > ± 50% change. The top 5 proteins by logFC are labeled. (d) Heatmap of secreted proteins exhibiting a sex difference at both ZT8 and ZT20. (e) For each secreted protein, its logFC in the secreted fraction is plotted with its logFC in the liver, to correlate changes in the two compartments. Pearson correlation, ****p<0.0001. Simple linear regression line shown. (f) Bubble plot of Gene Ontology enrichment (DAVID) of secreted proteins from d. Top 5 terms from each category are shown. Source data for (a) are provided as Supplementary Source Data.

**Supplementary Fig. 4.**
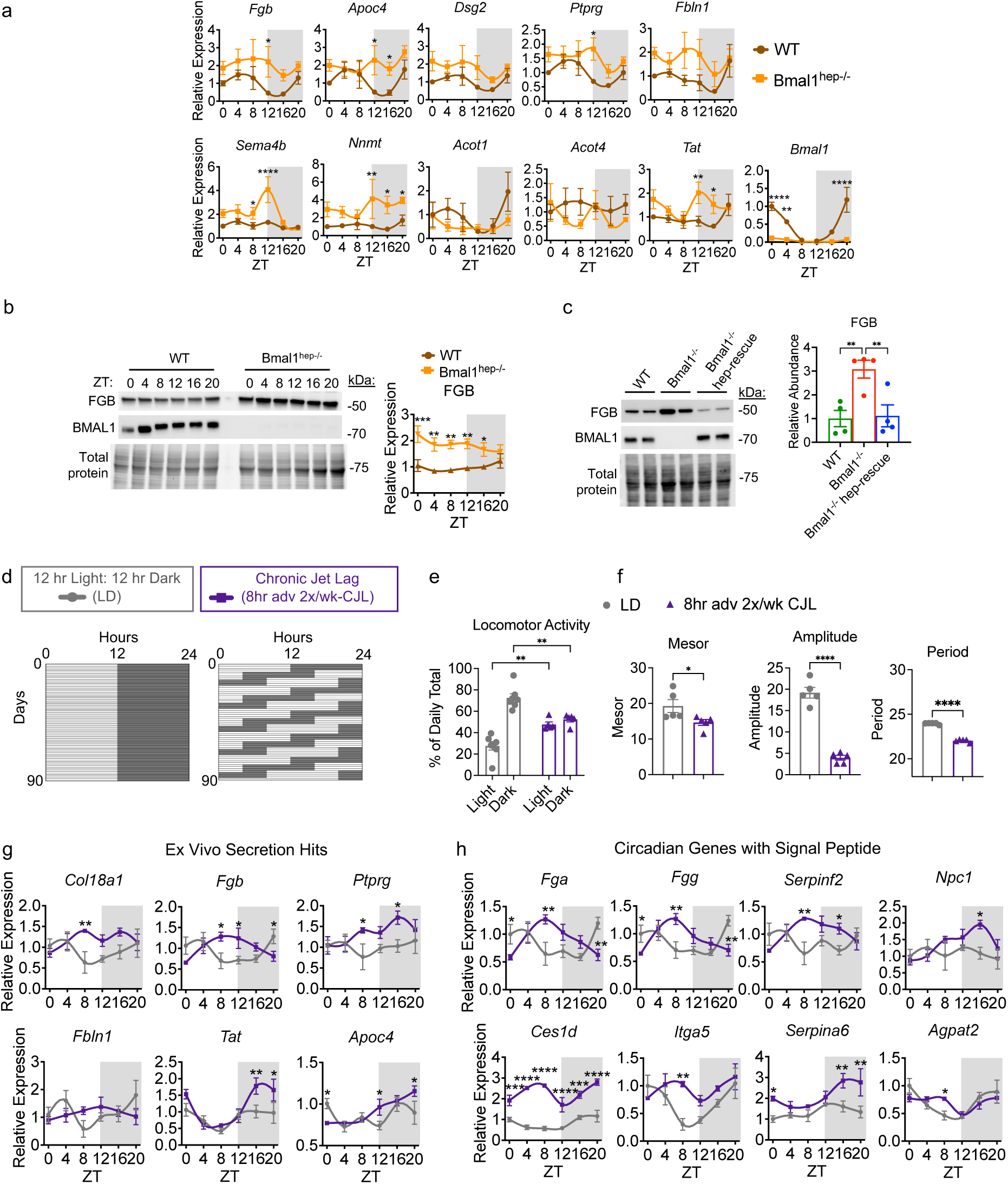
Data Related to Fig. 3. (a) Expression of time-dependent secreted proteins by qPCR in the liver at 6 diurnal timepoints. Two-way ANOVA, Fisher’s LSD post hoc test, *p<0.05, **p<0.01, ***p<0.001, ****p<0.0001. All groups n=4 except for *Apoc4, Dsg2, Ptprg, Sema4b, Nnmt, Acot1, and Tat* WT ZT0 and *Bmal1*^hep-/-^ ZT12 n=3, *Fgb* WT ZT4, 8, 16 and *Bmal1*^hep-/-^ ZT4, 12, 20 n=3, *Fbln1* WT ZT0 and *Bmal1*^hep-/-^ ZT4 n=3, *Acot4* and *Bmal1* WT ZT0 n=3. (b) Western blot from whole liver lysates collected at the indicated timepoints, replicates quantified to the right, Two-way ANOVA, Fisher’s LSD post hoc test, *p<0.05, **p<0.01, ***p<0.001. All groups n=4 except for WT ZT4, 8, 16 and *Bmal1*^hep-/-^ ZT4, 12, 20 n=3. (c) Western blot from whole liver lysates collected at ZT8, replicates quantified to the right, One-way ANOVA, Fisher’s LSD post hoc test, *p<0.05, **p<0.01, n=4. (d-h) Adult male C57BL/6J mice exposed to chronic jet lag (CJL), consisting of an 8-hour advance of the light schedule twice per week for 3 months from 4 to 16 weeks-of-age. (d) Schemes of normal light-dark (LD) and CJL light schedules. (e-f) Analysis of locomotor activity during the last 2 weeks of the paradigm. (e) Mixed effects model, Sidak’s post hoc test, **p<0.01, n=6. (f) Unpaired t test, *p<0.05, ****p<0.0001, n=5. (g-h) Expression of ex vivo secretion hits and known circadian genes with signal peptides by qPCR in the liver at 6 timepoints over the diurnal cycle. Two-way ANOVA, Fisher’s LSD post hoc test, *p<0.05, **p<0.01, ***p<0.001, ****p<0.0001, n=4. Source data for (a)-(c) and (e)-(h) are provided as Supplementary Source Data.

**Supplementary Fig. 5.**
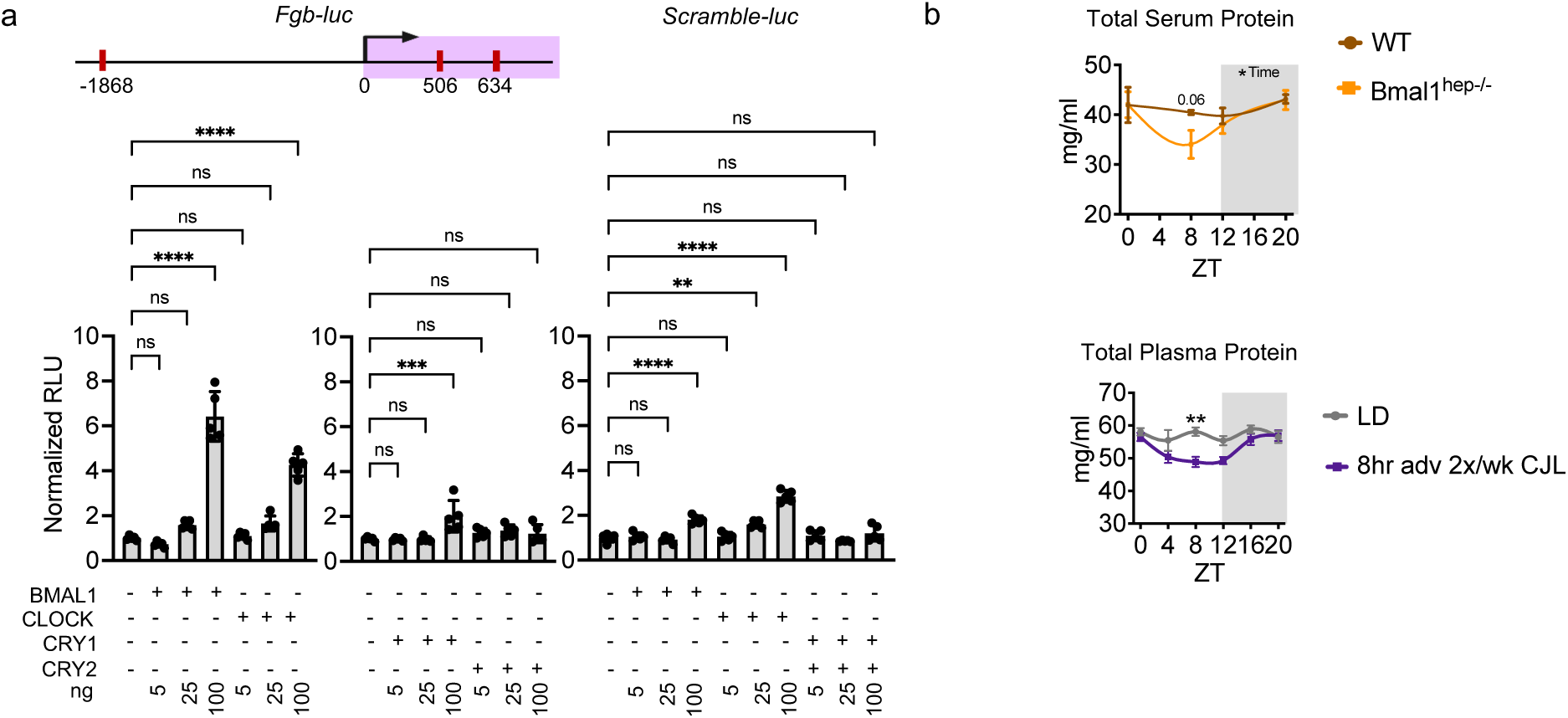
Data Related to Fig. 3. (a) AML12 hepatocytes transfected with the indicated luciferase (luc) reporter plasmids. One-way ANOVA, Tukey’s post hoc test, ***p<0.001, ****p<0.0001, n=5. (b) Top - Total protein concentration in serum. Two-way ANOVA, Sidak’s post hoc test, time *p=0.0549. All groups n=5 except for WT ZT8 n=4. Bottom - Total protein concentration in plasma. CJL – chronic jet lag. Two-way ANOVA, Sidak’s post hoc test, **p=0.0010. All groups n=5 except for LD ZT4 and ZT12 n=4. Source data for (a) and (b) are provided as Supplementary Source Data.

**Supplementary Fig. 6.**
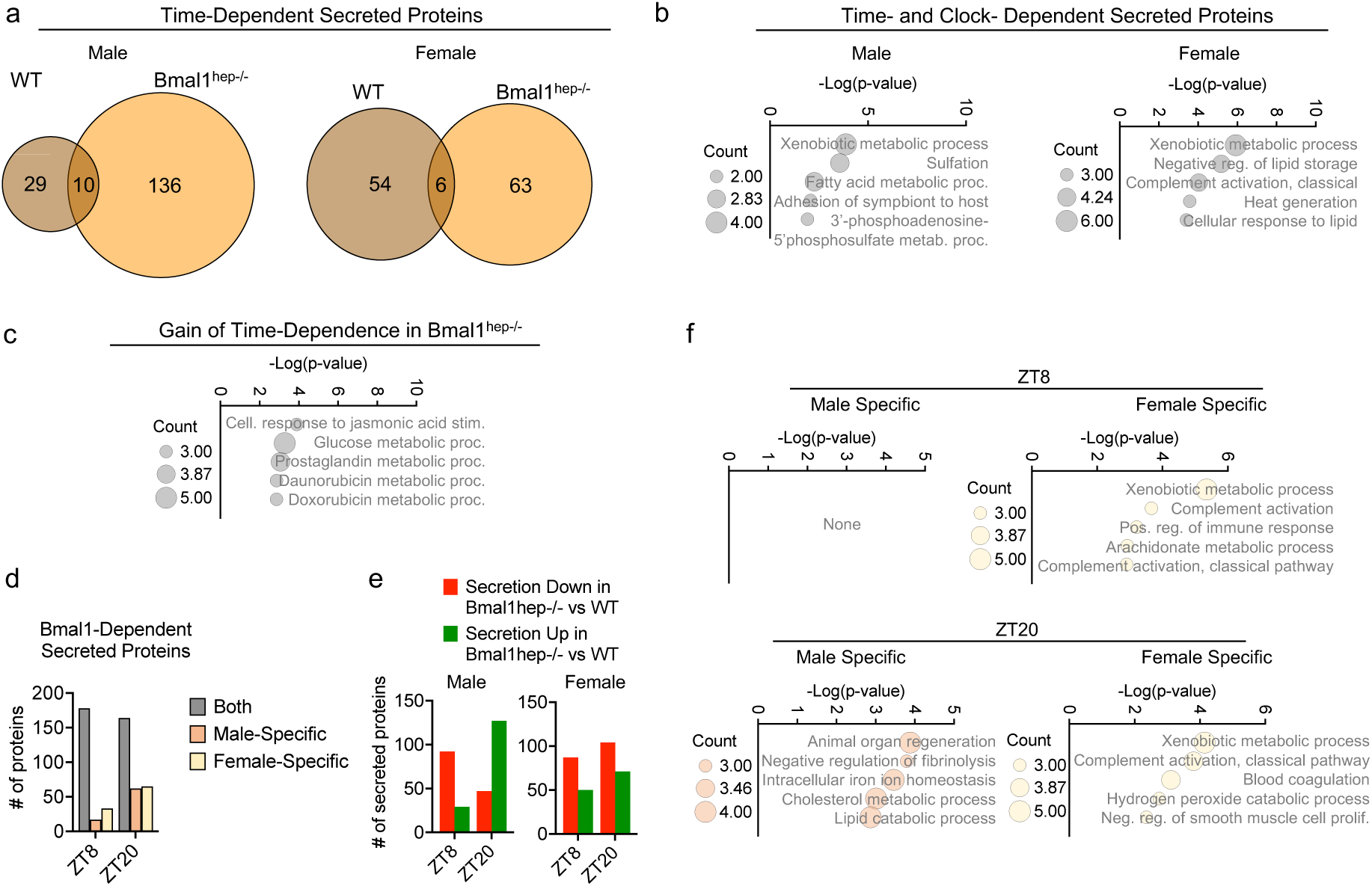
Data Related to Fig. 3. Analyses related to Fig. 3. (a) Analysis of time-dependent secreted proteins (Student’s t-test, FDR <0.1, > ± 50% change) in WT and *Bmal1*^hep-/-^mice according to sex. (b) Gene Ontology enrichment (Biological Process) analysis of time- and clock- dependent secreted proteins (signal peptide and unconventional secretion) according to sex. (c) Gene Ontology enrichment (Biological Process) analysis of secreted proteins (signal peptide and unconventional secretion) that gained time-dependence in *Bmal1*^hep-/-^. (d) Breakdown of *Bmal1*-dependent secreted proteins according to sex. (e) Direction of change for *Bmal1*-dependent secreted proteins according to sex. (f) Gene Ontology enrichment (Biological Process) analysis of all *Bmal1*-dependent secreted proteins at each time point according to sex.

**Supplementary Fig. 7.**
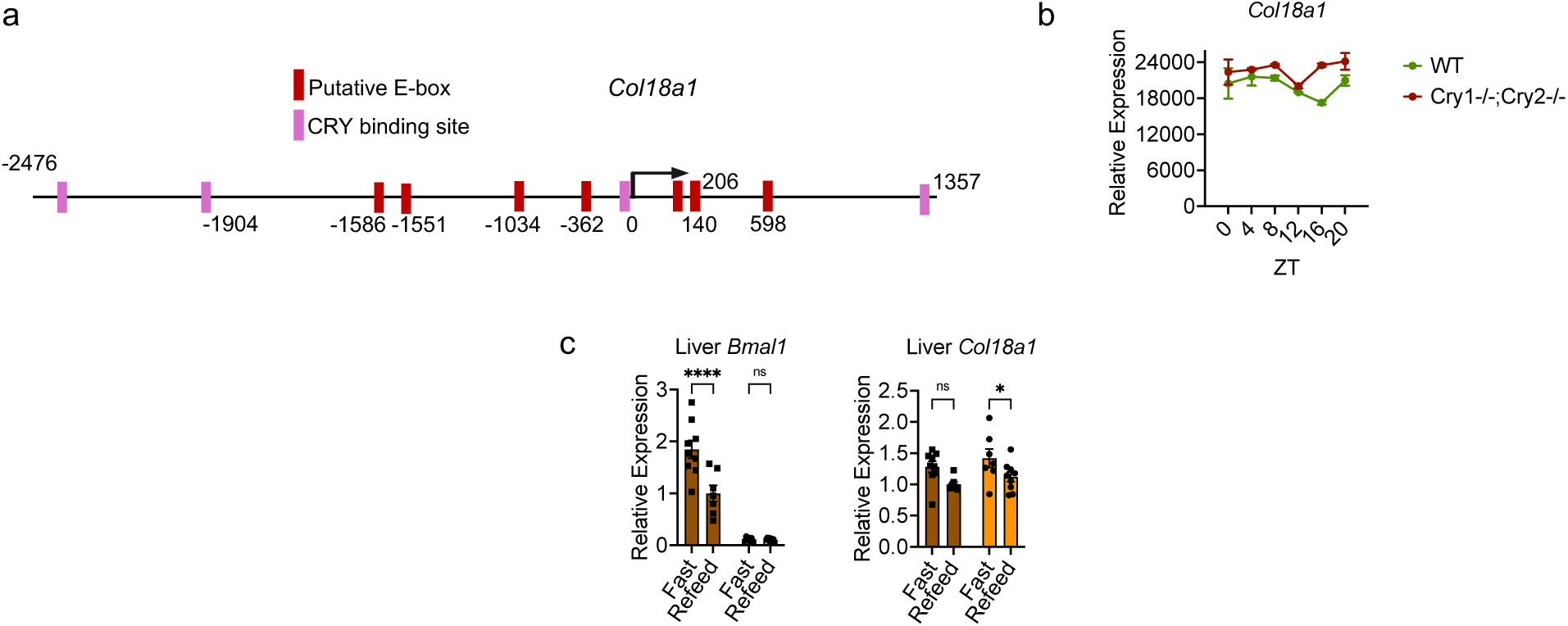
Data Related to Fig. 4. (a) Location of putative E-boxes according to the Eukaryotic Promoter Database, JASPAR core 2018 vertebrates, p<0.001. Location of CRY binding sites (CRY1 or CRY2, at CT 0, 4, 8, 12, 16, or 20) from Koike et al., Science, 2012. (b) Liver RNA-sequencing data from Weger et al., PNAS, 2021. (c) qPCR in liver, Two-way ANOVA, Fisher’s LSD, *p<0.0398, ****p=<0.0001, WT refeed male and *Bmal1*^hep-/-^ fast male n=3, WT refeed female, WT fast female, *Bmal1*^hep-/-^ refeed female, and *Bmal1*^hep-/-^ fast female n=4, WT fast male and *Bmal1*^hep-/-^ refeed male n=5. Source data for (b) and (c) are provided as Supplementary Source Data.

**Supplementary Fig. 8.**
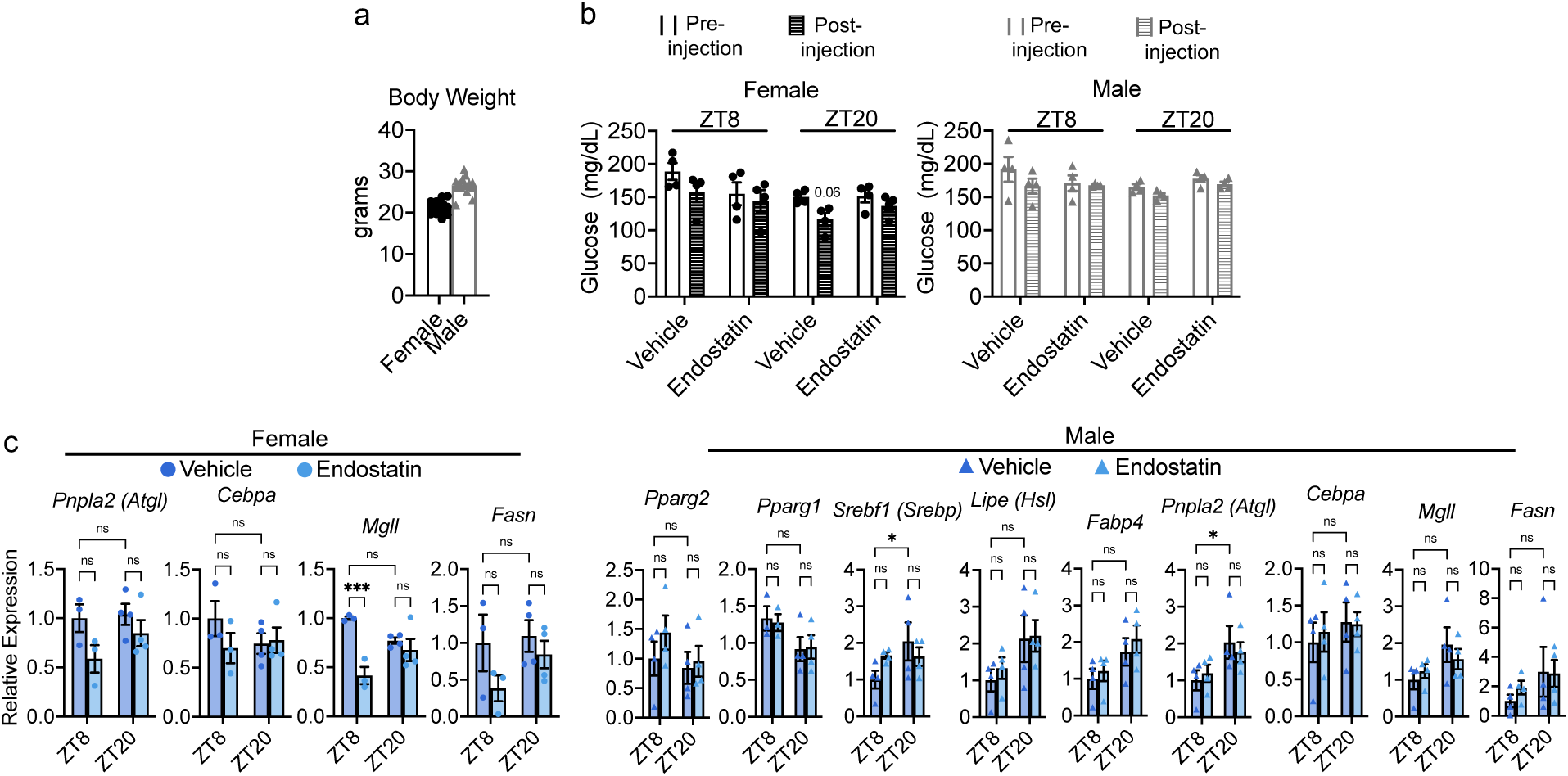
Data Related to Fig. 5. (a) Body weight at the time of endostatin injection. (b) Blood glucose values pre- and post-endostatin injection, n=4. (c) qPCR in epididymal white adipose tissue (eWAT). Two-way ANOVA, Fisher’s LSD post hoc test, *Mgll* ***p=0.0006, *Srebf1* *p=0.0378, *Pnpla2* *p=0.0386, n=3-4 (n-value for each group shown within graph). Source data for (a)-(c) are provided as Supplementary Source Data. Supplementary Source Data:

## Notes

### Competing Interest Statement

The authors have declared no competing interest.

### Summary of Updates

This is a revised version of the manuscript, in response to rounds of reviewer comments.

